# Inter-population connectivity of southern elephant seals and the likely intra-species transmission pathways of high pathogenicity avian influenza

**DOI:** 10.64898/2026.07.07.737127

**Authors:** Clive R. McMahon, Mark A. Hindell, Robert G. Harcourt, Isobel Lerpiniere, Ian Jonsen, Christophe Guinet, Rupert Woods, Marthan N. Bester, Jane Younger, Nicholas M. Fountain Jones, Tristan Burgess

## Abstract

High Pathogenicity Avian Influenza (HPAI) H5N1 clade 2.3.4.4b has spread beyond birds to affect seals across the Southern Ocean and sub-Antarctic region, with southern elephant seals (*Mirounga leonina*) particularly devastated. The virus, likely introduced via spillover from infected migratory birds, has killed tens of thousands of adult seals and pups throughout most of their range, though Macquarie Island remains unaffected so far.

We used twenty years of elephant seal movement data from the southern Indian and Pacific oceans to assess whether seal-to-seal transmission could spread HPAI H5N1 between breeding colonies, despite the vast distances separating them (Marion Island, Iles Crozet, Iles Kerguelen, and Macquarie Island). There was substantial overlap in seals’ at-sea distributions during their winter post-moult trips, when seals travel for weeks at average speeds of 3.5 km/h.

Two transmission pathways were examined: (1) terrestrial “stepping stone” routes, where infected seals could pass the virus between colonies during short intervals to remain infectious were feasible from Marion Island to Kerguelen but not from Kerguelen to Macquarie Island; and (2) at-sea encounters between seals, which occurred frequently enough to enable transmission.

The findings suggest that once established at Macquarie Island, the virus could potentially spread further to New Zealand’s sub-Antarctic islands and mainland New Zealand. While seal-to-seal transmission appears possible, we conclude this is unlikely. Nonetheless, understanding at-sea contact rates enhances knowledge of H5N1 epidemiology and demonstrates the value of combining long-term population monitoring with movement data to understand wildlife disease dynamics.

## Introduction

Since 2021, High Pathogenicity Avian Influenza H5N1 clade 2.3.4.4b has spread widely across the globe, spilling over into mammals from its original avian hosts (Leguia *et al*. 2023; Xie *et al*. 2023; Bi *et al*. 2024; Uhart & Vanstreels 2025; Xie *et al*. 2025; Knief *et al*. 2026). This spillover includes seals and probably occurred through direct bird-to-mammal contact at dense breeding and moulting colonies and/or through environmental exposure. In the Southern Hemisphere seal breeding and moulting sites are often near large dense penguin breeding and moulting areas^7.^ with the seals’ seasonal and geographic locations often overlapping with that of potentially infected avian host species increasing the risk of virus transmission. Marine mammals have been disproportionally and negatively affected by this spillover with some notable examples in the Southern Ocean, with pup mortality rates up to 97% and major declines in colony populations reported (Campagna *et al*. 2023; Bamford *et al*. 2025; Clessin *et al*. 2025; Ashley *et al*. 2026; Clessin *et al*. 2026). The Southern Ocean and its subantarctic islands host dense seabird colonies and seal populations (including southern elephant seals, Antarctic & subantarctic fur seals) that could encounter viruses shed by infected avian hosts, either directly or via environmental media. Viral phylogenetic reconstructions of HPAI H5N1 detected on the sub-Antarctic island of Crozet and Kerguelen provide strong evidence for these introductions having distant and multiple independent sources (Campagna *et al*. 2023; Bamford *et al*. 2025; Clessin *et al*. 2025; Ashley *et al*. 2026; Clessin *et al*. 2026). These introductions into the Crozet (5,600 km) and Kerguelen (6,600 km) archipelagos and their phylogenetic linkages to South Georgia underscore the complex, trans-regional pan-oceanic dispersal dynamics of the virus. Aside from the Antarctic Peninsula, the virus has not been reported on mainland Antarctica. Macquarie Island in the southern Pacific Ocean (5755 km from Kerguelen) is the only major sub-Antarctic seabird and seal colony yet to be infected (as of July 2026). This highlights the urgent need to understand more comprehensively (1) potential transmission pathways, 2) intra- and inter-population connectivity, (3) HPAI H5N1 presence, occurrence and prevalence across the Southern Ocean.

Currently, the view of viral transfer between Antarctic and sub-Antarctic mammal populations, is that seabirds are most likely to move the virus from one population to another (Kuiken *et al*. 2026). The presumed transport hosts are predatory or scavenging species (such as skuas or giant petrels) given these species are at high risk of exposure to HPAI H5N1 and are thought to be the most likely to facilitate the jump between populations while remaining alive and infectious(Clessin *et al*. 2025). Alternative transmission models that include staged transmission via a series of stepping stones (multiple individuals), and perhaps involving multiple species have been postulated, but are difficult to demonstrate quantitatively due to the paucity of tracking data(Riaz *et al*. 2024). The distances between Antarctic breeding populations can be considerable, nonetheless the virus has successfully reached Iles Kerguelen, 6,600 km from the known source at South Georgia(Clessin *et al*. 2025). Because most of the affected species, such as southern elephant seals, fur seals and various penguin species, spend most of their lives at sea, only aggregating on land for annual breeding or moulting periods it is useful to consider alternative transmission pathways/models. One possible transmission model is at-sea seal-to-seal transmission of the virus during the winter pelagic phase of their annual lifecycle. Our ability to assess this possibility relies on good tracking data which permits quantitative understanding of habitat use and informs key parameters such as encounter rates. These data are not available for most species.

The notable exception is the southern elephant seal, which is also one of the Antarctic species most severely impacted by HPAI H5N1 (Campagna *et al*. 2023; Bamford *et al*. 2025; McInnes *et al*. 2026). Southern elephant seals typically make two long distance migrations each year, spending up to 8 months at sea and traveling up to 6,000 km from their breeding and moulting sites (Hindell & McMahon 2000; Hindell *et al*. 2016; Hindell *et al*. 2021). This means that it is possible for elephant seals to move between all of the major breeding sites, potentially providing an alternative transmission pathway. However, the pattern is complex with different age and sex classes of seals having different migration patterns and timing, and most animals returning to the same breeding sites each year (Hindell & Little 1988). Elephant seals have been the subject of a plethora of tracking studies since the 1990s which have resulted in a comprehensive understanding of the species’ at-sea distribution and movements (Hindell *et al*. 2020) making it one of the best studied species in this regard globally. For example, over 800 elephant seals of various age and sex classes have been tagged at Iles Kerguelen since 2004 as part of a long-term integrated biology and ocean physics study(McMahon *et al*. 2025), and all of the other major populations have also been studied to some extent(Hindell *et al*. 2016). Here we use a detailed compilation of elephant seal tracking data to explore the likelihood of seal-to-seal transmission of HPAI H5N1. Here we use a detailed compilation of elephant seal tracking data to explore the likelihood of seal-to-seal transmission of HPAI H5N1 in the southern Indian and Pacific Oceans.

We consider two, non-mutually exclusive, ways for this to happen. The first is the virus moving between island populations via a series of land-based transmission events at terrestrial haulout sites where the seals aggregate outside the breeding season to moult and rest. There are a number of these sites in East Antarctica such as the Vestfold Hills, the Windmill Islands and Commonwealth Bay which could theoretically act as stepping stones for the virus to move from Iles Kerguelen to Macquarie Island (Bester 1988; van den Hoff, Davies & Burton 2003; Bester *et al*. 2020; Chua *et al*. 2022) (the only major elephant seal population not so far affected by HPAI H5N1).

Alternatively, the virus could be transmitted between seals while at sea. At present there is only indirect evidence of the occurrence of such interactions, for example, it has been hypothesised that some mating is likely to occur at sea(de Bruyn *et al*. 2011). Also, it has been hypothesised that there may be a pool of diseased animals at sea during the winter months which serve as the source of viral re-introduction to South Georgia in the subsequent breeding season, permitting the observed year-to-year persistence(Bamford *et al*. 2025). As this is a period of 6 to 8 months, far longer than the infectious period of any individual, such a pool of infected animals could only realistically persist over winter if the virus is transmitted between individuals at sea. The population size for some breeding colonies is considerable (approximately 110,000 breeding females for South Georgia (Bamford *et al*. 2025) and 99,247 breeding females for IIes Kerguelen (Laborie *et al*. 2023) prior to the current outbreak), which combined with the observation that the seals tend to aggregate where prey is abundant *i.e.* in areas of ecological importance(Hindell *et al*. 2020), this raises the possibility of seals from different breeding colonies encountering each other during their time at sea. The encounter rates may be enhanced in the pack-ice where seals potentially haul-out to rest (de Bruyn *et al*. 2011; Hindell *et al*. 2020; Laborie *et al*. 2023; Bamford *et al*. 2025).

We use the combined tracking dataset from three breeding colonies in the South Indian and Pacific Oceans to explore the possibility that seals could carry HPAI H5N1 from Marion Island and/or Iles Kerguelen (both of which have HPAI H5N1 in their elephant seal populations) to Macquarie Island, which was free of any signs of HPAI H5N1 in the 2025/2026 summer period. We use the data to calculate the at-sea rates of transit for the seals, as this may be a key factor in how far an infectious seal can move the virus. We then explore (i) the possibility of the virus moving between islands via terrestrial haulout sites, by quantifying the at-sea travel rates of the seals and describing potential incubation and infectious periods of the virus that would be necessary for the transmission to be possible, and (ii) the possibility the virus is transmitted between seals that encounter each other at sea, by quantifying the at-sea encounter rates of tracked seals both within and between the three colonies. While HPAI H5N1 outbreaks in the Southern Ocean are currently thought to be facilitated by movement of scavenging bird species (Uhart *et al*. 2024; Clessin *et al*. 2025; Clessin *et al*. 2026) as mammal-mammal transmission increases in frequency and the virus accrues adaptions to mammalian hosts, the probability of mammals to aid in transporting this virus increases. Movement data such as these provide a powerful lens through which to study possible transmission routes of viruses, quantify contact rates, identify possible transmission hotspots, track pathogen spread, and ultimately predict outbreaks, anticipate spillover events and design better management strategies.

## Methods

### Tracking data

We have focused on three of the four major elephant seal breeding sites in the Southern Indian and Pacific Ocean. We focused on two populations where tracking data is available and HPAI H5N1 virus was reported in the austral summer of 2024 - Marion Island and Iles Kerguelen(Clessin *et al*. 2025) - and on Macquarie Island, 2,500 km to the east of Iles Kerguelen, which has yet to record either confirmed or suspect cases. As part of a long-term monitoring study, starting in 2004, we have attached conductivity temperature depth satellite relayed data loggers (CTD-SRDLs; Sea Mammal Research Unit, University of St. Andrews, UK) to a total of 791 post-moult female and sub-adult male southern elephant seals (McMahon *et al*. 2025): 586 at Iles Kerguelen, 121 at Macquarie Island and 84 at Marion Is. (Figure 1). The capture, handling, sedation and attachment procedures are described fully elsewhere(McMahon *et al*. 2000; Field *et al*. 2002; McMahon *et al*. 2008; Field *et al*. 2012).

**Figure 1.**
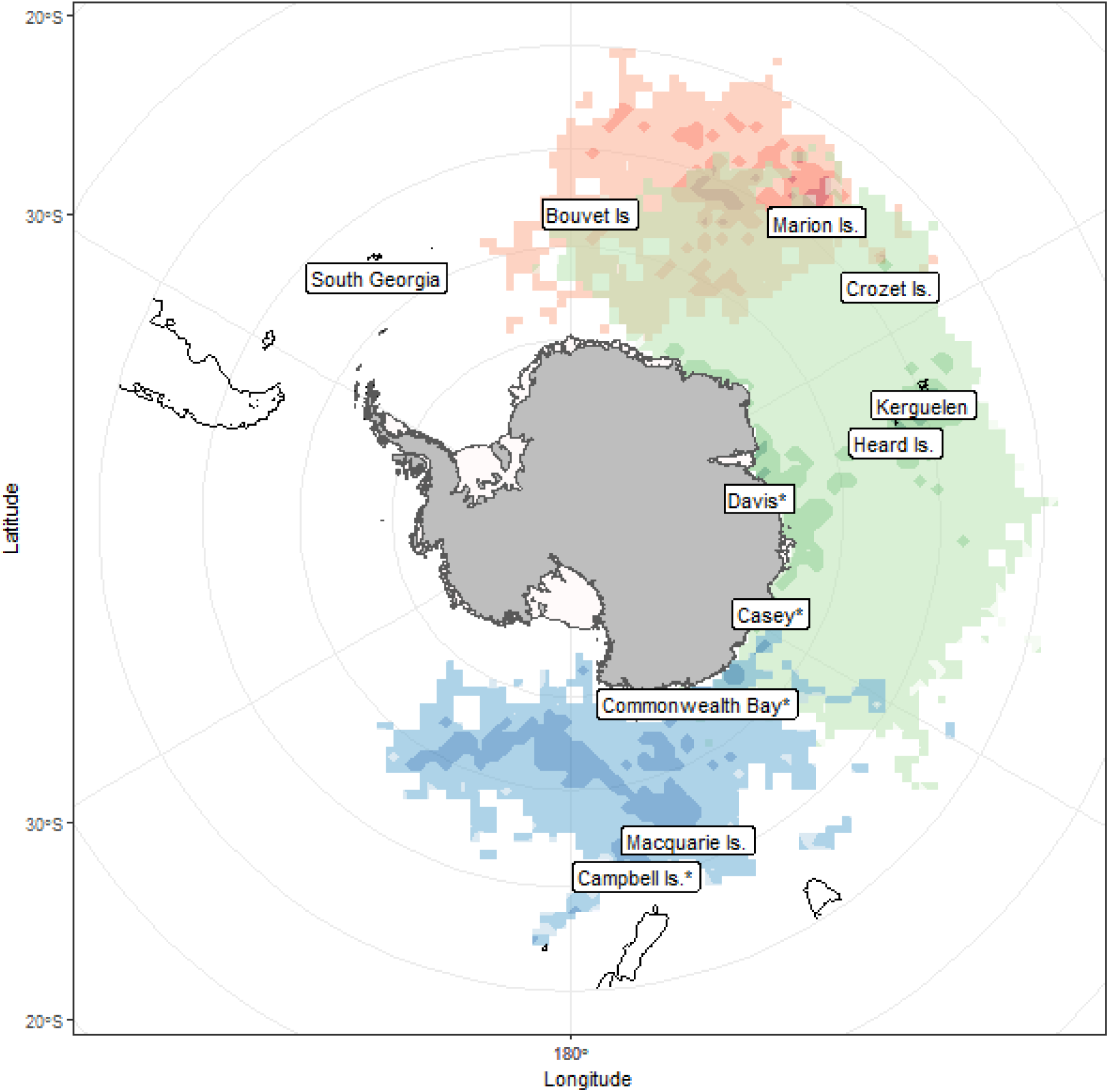
Map of Antarctica and the Southern Ocean showing the at-sea utilisation distributions of southern elephant seals from the breeding colonies of Marion Island (n = 84, red), Iles Kerguelen (n = 586, green) and Macquarie Island (n = 121, blue). Darker shades indicate the 50% UD (enclosing the cells containing the top 50% of the total seal days for that site). The asterisk (*) denotes haul-out only sites.

The real-time at-sea Doppler-derived locations provided by Argos (Argos 2016) were filtered using a random walk state-space model (SSM) in the R package *aniMotum* (Jonsen *et al*. 2023) after removing locations before the start and end of the foraging trip and also removing seals with fewer than five days of data. We used a 6-hour timestep (estimating a location at 00:00, 06:00, 12:00 and 18:00 each day) with a maximum rate of travel of 4 m^sec^, to obtain an estimated location (latitude and longitude) with its associated uncertainty.

### Rate of travel at sea for southern elephant seals

A key parameter in identifying possible transmission rates is the distance seals travel within a given period. We calculated the horizontal swimming speed of the seals based on the 6-hourly locations estimated during the path analysis step. Elephant seals make migrations lasting from 3 to 8 months, depending on their age and sex class. The nature of these trips varies widely among individuals but, can broadly be characterised as having an outward phase lasting days to weeks, when the seals undertake rapid, directed travel, followed by a period of less rapid and more meandering travel (known as area restricted search (ARS)), followed by a period of rapid and directed travel back to the colony.

To determine the most likely rates of travel when the seals are potentially infectious (assuming they become infected at their breeding or moulting sites), we isolated the initial outward phase of each seals’ migration (Figure 2a) and calculated the median speed between all of the 6-hourly locations along that section of the track. We defined the outward phase as the distinctly linear phase of the distance from the island over time, marking the start of the trip as when the seal left the island and identifying the end of the outward phase when the seal started its first period of ARS (Figure 2b). We then compiled all the median speeds from the 791 seals and calculated the overall median speed and the upper 90^th^ percentile of the distribution to provide two alternative rates of travel for the subsequent analyses (Figure 2c). Because tracking data from HPAI-infected elephant seals are not available, these movement rates are derived from seals that were not known to be clinically affected. We therefore interpret these as plausible rates for infected seals that remain subclinical, pre-symptomatic, mildly affected, or otherwise sufficiently mobile to undertake directed travel. Clinically affected seals may travel more slowly, less directly, or not complete sustained outward movements, meaning that the median and upper-percentile values used here may overestimate transmission distances for severely ill individuals. We regard this is likely to be realistic given that such mildly affected animals, to the extent that they exist, represent the best candidates for long-distance viral transmission by seals.

**Figure 2.**
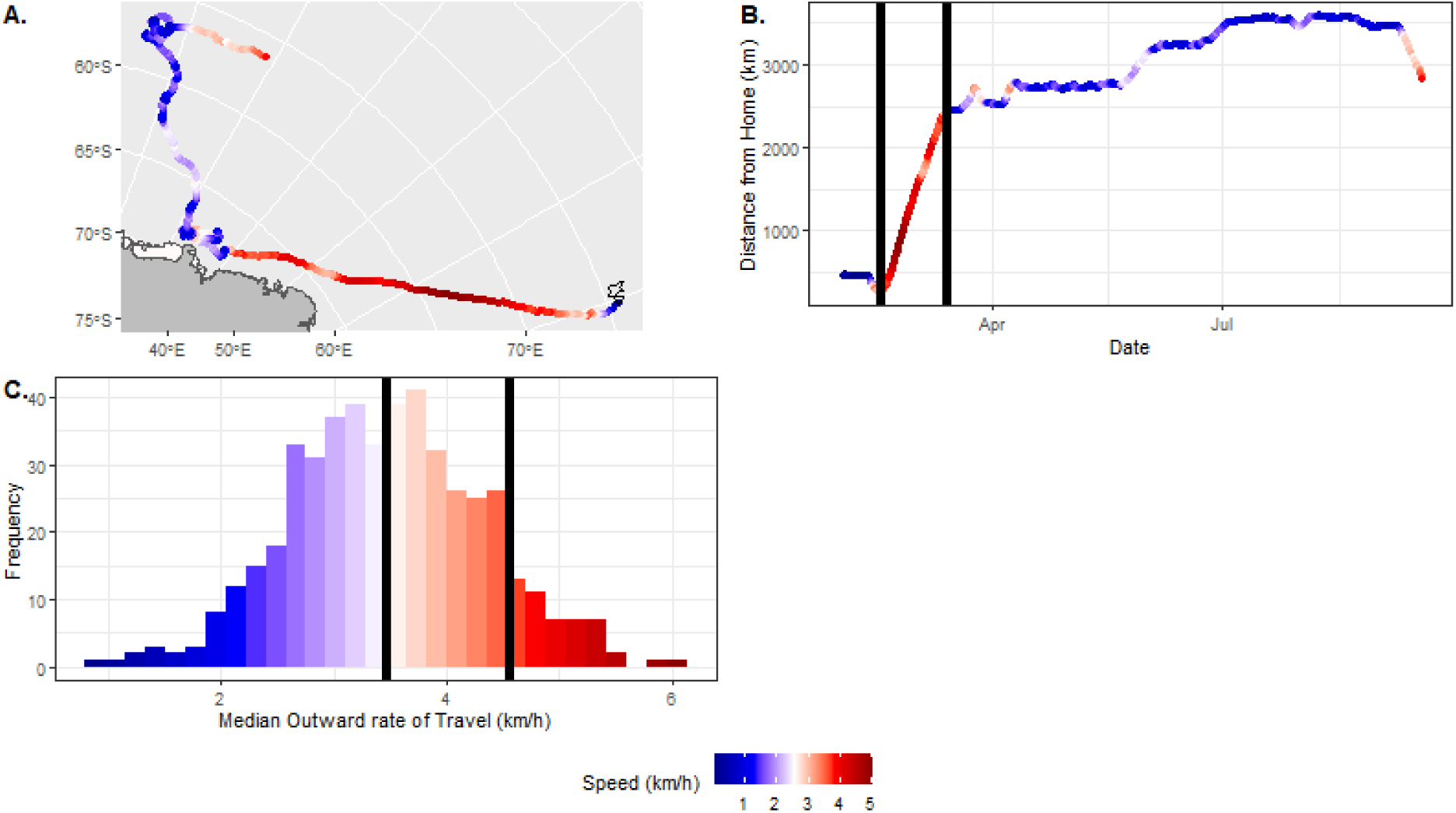
Analysis of rates of travel during the outward leg of southern elephant seal trips to sea. (A.) Example of a track (seal ID = ct105-14f-13) deployed at Iles Kerguelen in February 2014), with the six hourly locations colour-coded by the speed (km/h) between successive points. (B.) the distance from Kerguelen of each 6 hourly location throughout the trip (here the tag stopped on 18 September 2014, before the seal returned to Iles Kerguelen). The two vertical lines denote the period of outward travel, which lasted for four weeks during which the seal had a median speed of 3.9 km/h. (C.) The frequency distribution of median outward speeds for each of the 791 seals in the study. The first vertical line marks the 50^th^ percentile of the distribution (3.46 km/h) and the second line indicates the 90^th^ percentile of the distribution (4.55 km/h).

### Possible terrestrial transmission routes

To quantify the likelihood of elephant seal-to-elephant seal transmission of the virus between the colonies and haulout sites we combined information on at-sea transit rates (Figure 2) with the distances between known haulout sites and an assumed latent period of 4 days. This allowed us to determine the number of days a seal would need to be infectious for the virus to be transmitted between each known haul-out or breeding site. We then assessed the likelihood of a virus being transmitted along each step based on this infectious period, identifying those with a required infectious period of less than 14 days as potential steps (*i.e.* representing those where transmission is plausible). We based the possible latent and infectious periods on a review of the literature on mammalian latent and infectious periods for influenza A viruses. Because the frequency of sub-clinical infection and movement impairment are unquantified for HPAI in free-ranging seals, we must assume that a proportion of infections must remain subclinical or to retain sufficient mobility for transmission during the infectious period for this transmission pathway to be possible. This assumption is epidemiologically important because severe mortality has been documented in elephant seals, including the large Peninsula Valdés outbreak(Uhart *et al*. 2024), but the frequency of subclinical infection, pre-morbid movement, and effective infectiousness before death remain unknown. These parameters are likely to vary with age class, exposure route, viral genotype, and potentially ongoing adaptation associated with mammal-to-mammal transmission. Though rare, asymptomatic infection has been reported to occur at low levels even in viral infections such as Ebola Virus Disease which are traditionally viewed as having very high morbidity (Glynn *et al*. 2017).

We also assume that the seal movements occur in the summer and autumn when both males and females are in their post-moult migrations and the sea-ice is at its minimum, allowing access to continental Antarctic haul-out sites. Although male and female at-sea ranges differ somewhat (with males more likely to go onto the Antarctic continental shelf) we have not modelled them separately as both sexes do use the shelf to some extent.

### Possible at-sea transmission routes

Two lines of evidence suggest that seals may encounter each other while at sea, creating opportunity to pass the virus from one to the other: (i) southern elephant seals do not move about the ocean at random, but have preferred regions that they use more than others (Figure 1) and (ii) There are large populations of southern elephant seals, particularly in the Kerguelen population (350,000 in total), so there are likely to be relatively high densities of animals concentrated in these preferred regions, increasing the prospects of two or more seals being in the same place at the same time.

We used at-sea seal tracking data to quantify the probability of at-sea encounters. We initially did this for the observed times (*e.g.* two tracked seals present in the area at the same time). We identified each of the 6-hour time steps that contained more than one seal and then calculated all the pairwise distances between those seals (using the *distm* function in *geosphere* R package). We refined this further by then defining two seals as potentially being in the same place simultaneously as when the SSM-estimated location error ellipses of a pairwise comparison overlapped (to any extent).

However, the observed data are limited due to the seals being sampled over a 20-year period, with no more than 40 seals tracked in any one year, which will greatly underestimate the actual occurrence of at-sea encounters. We therefore conducted an analysis to estimate the likely encounter rate at a scale more akin to the overall population level. This was simply done by assuming that all the seals were tracked in the same year, effectively creating a much larger annual sample (n = 791), then repeating the calculation of at-sea encounters describe above. Importantly, we restricted this analysis to encounters that were > 200 km from the breeding colonies to avoid biasing the analysis with the high concentration of encounters that occurred close to the island (this is because all the seals pass through these regions on their way to and from the islands). We then calculated: (i) the number of other seals that each seal encountered during its time at sea; (ii) the number of times each of the encountered seals is encountered (a proxy of the time that the seal spends with each of the other seals that it encounters); (iii) the duration between successive encounters and (iv) the distance between each encounter.

## Results

The tracked seals from all three breeding sites (Marion Is, Iles Kerguelen and Macquarie Is) dispersed widely during their migrations (Figure 1), with some individuals travelling several thousand kilometres. Importantly, the regions used during these migrations overlapped between the colonies. More than half of the areas used by Marion Island seals were also visited by seals from Iles Kerguelen. Being further apart, the overlap between the Iles Kerguelen and Macquarie seals was less, but not zero. Seals from all colonies visited the Antarctic continent, but there are no known haulout sites in the area directly south of Marion Island, so there was limited opportunity for the seals from Marion Is and Iles Kerguelen to interact on the Antarctic continent. However, they could do so at Crozet Is, approximately mid-way between Marion Is and Iles Kerguelen. The seals from Iles Kerguelen and Macquarie Is have three known Antarctic haulout sites, the Vestfold Hills (near Davis Station) and the Windmill Is, (near Casey Station) and Commonwealth Bay (Figure 1), presenting a possible series of stepping stones between the two colonies.

### Daily rate of travel

The individual speeds of seals during their outward phase ranged considerably, from less than 2km/h to over 6 km/h (Figure 2c). The median value was 3.5 km/h and the 90^th^ percentile (speeds achieved or exceeded by the fastest 10% of the population) was 4.6 km/h. These values were used in the following assessment of terrestrial transmission potential of the virus.

### Possible terrestrial routes of transmission

The shortest distance between a known HPAI-affected site and Macquarie Island is 5,247 km (Heard Island to Macquarie) (Table 1), which is most likely too far for a single infected seal to carry the virus - even travelling at the 90^th^ percentile of known transit speeds (4.6 km/h) it would take 46.5 days for seals to make the journey uninterrupted. This suggests that if seals were to carry the virus between these sites, it would have to be via a series of intermediate steps involving either terrestrial haulout sites or at-sea encounters. Assuming a continued west to east spread pattern of the virus, a plausible terrestrial route could be: 1. Kerguelen to Heard Is (525 km taking 4.6 d), 2. Heard Is to Davis (1,739 km, taking 15.4 d), 3. Davis to Casey (1,397 km, taking 12.4 d), 4. Casey to Commonwealth Bay (1,402 km, taking 12.4 d) and finally 6. Commonwealth Bay to Macquarie Is (1614 km, taking 14.3 d) (Figure 3). Assuming a four-day latent period for the virus, a seal would need to remain infectious for 11.4 days for the virus to be transported from Heard Is to Davis, the longest leg in this possible transmission route. Estimates of viral shedding duration or infectious period are not currently available for HPAI H5N1 in pinnipeds (Uhart *et al*. 2024; Uhart & Vanstreels 2025). Although this exceeds the typical infectious period in mammals, prolonged human influenza A (H1N1) viral shedding exceeding 14 days has been reported in some humans (Fielding *et al*. 2014), particularly in more severe infections (though including some mild cases), suggesting that transport over this route is not biologically implausible. In the absence of host species-and strain-specific estimates for HPAI H5N1 infectious period in southern elephant seals, this route should therefore be considered biologically plausible but currently weakly constrained.

**Figure 3.**
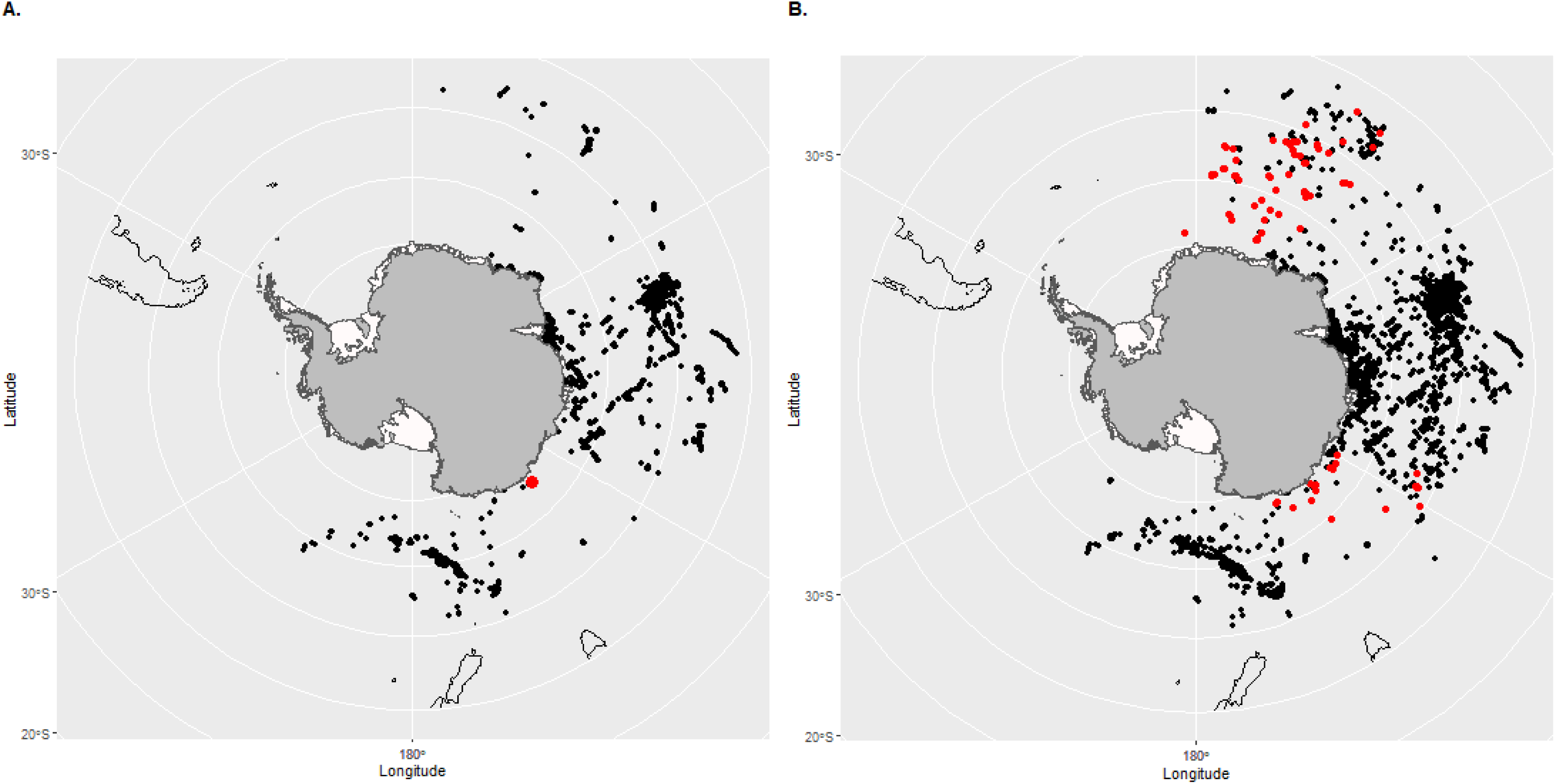
Two maps of potential at-sea encounters of elephant seals, defined as where the uncertainty ellipses of a pair of locations from the same time overlap. Black dots indicate locations where the pair of seals were from the same breeding colony and red indicates where the pair of seals are from different breeding colonies. A. The real-time data, where the proximity of the pairs of seals are based on year, month and day and time. Of a total of 33,285 potential at-sea meetings only one (highlighted by the red dot) was from two separate colonies (in this case Macquarie Island and Iles Kerguelen 2005-05-01 12:00:00). B. Results of a simple population level analysis where we assumed that all of the sample of 786 seals were all deployed in the same year. In this case there were 200,661 potential at-sea meetings between pairs of seals, 116 of which were between seals from different breeding colonies (red dots).

**Table 1.** Matrix of distances (km) between the known breeding and haul out sites (*) for elephant seals in the southern Indian and Pacific Oceans. The lower diagonal is the straight-line distance between each of the sites (km). The upper diagonal matrix is the number of days that a seal would need to be infectious in order for the virus to arrive there from the starting location. This was calculated assuming a rate of travel of 4.7 km/h (Figure 1) and an initial incubation period of 4 days. It also assumes the seals leaves on the day it becomes infected and travels directly to new site. Those coloured in yellow are sites requiring an infectious period of less than 14 days. This would then provide a possible (but perhaps unlikely) series of stepping stones between Marion Is and Macquarie Is, from where seals could take it to New Zealand.

|  | South Georgia | Bouvet Is | Marion Is. | Crozet Is. | Kerguelen | Heard Is. | Davis* | Casey* | Commonwealth Bay* | Macquarie Is. | Auckland Is.* | Campbell Is.* | Antipodes Is.* |
| --- | --- | --- | --- | --- | --- | --- | --- | --- | --- | --- | --- | --- | --- |
| South Georgia |  | 19 | 41 | 48 | 54 | 53 | 43 | 52 | 54 | 65 | 68 | 66 | 67 |
| Bouvet Is | 2560 |  | 18 | 27 | 35 | 35 | 31 | 43 | 50 | 64 | 69 | 67 | 71 |
| Marion Is. | 5069 | 2536 |  | 5 | 17 | 19 | 25 | 37 | 48 | 62 | 68 | 67 | 73 |
| Crozet Is. | 5914 | 3468 | 1065 |  | 8 | 11 | 22 | 32 | 44 | 57 | 63 | 63 | 69 |
| Kerguelen | 6590 | 4396 | 2335 | 1343 |  | 1 | 16 | 23 | 35 | 47 | 53 | 53 | 59 |
| Heard Is. | 6460 | 4424 | 2621 | 1731 | 525 |  | 11 | 18 | 31 | 43 | 48 | 49 | 55 |
| Davis* | 5351 | 3958 | 3285 | 2879 | 2205 | 1739 |  | 8 | 19 | 33 | 39 | 38 | 44 |
| Casey* | 6323 | 5288 | 4611 | 4037 | 3011 | 2486 | 1397 |  | 8 | 21 | 27 | 27 | 33 |
| Commonwealth Bay* | 6534 | 6111 | 5894 | 5411 | 4418 | 3893 | 2609 | 1407 |  | 10 | 16 | 15 | 21 |
| Macquarie Is. | 7817 | 7692 | 7456 | 6881 | 5755 | 5247 | 4183 | 2857 | 1614 |  | 2 | 2 | 9 |
| Auckland Is.* | 8147 | 8226 | 8099 | 7536 | 6408 | 5901 | 4819 | 3508 | 2224 | 655 |  | 1 | 4 |
| Campbell Is.* | 7895 | 8058 | 8043 | 7525 | 6437 | 5923 | 4758 | 3488 | 2150 | 718 | 294 |  | 3 |
| Antipodes Is.* | 7987 | 8441 | 8650 | 8191 | 7143 | 6624 | 5375 | 4163 | 2782 | 1462 | 910 | 744 |  |

This transmission route is unlikely to involve adult females as they (1) rarely visit the terrestrial haulouts and (2) they are at sea for too long (∼ 8 months post-moult). Sub-adult males, often make two-to-three-month trips to sea punctuated by mid-year haul outs back at subantarctic colonies which typically occur in the first 6 months of the year (Hindell & Burton 1988; McMahon, Burton & Bester 1999). For the transmission route described above to be viable, an infected sub-adult male seal would need to leave Kerguelen in late December/early Jan, take 5 days to travel to Heard Is, where it then infects another seal. The Heard Island seal could leave in mid-January and spend approximately 16 days travelling to Davis, arriving in late January. There it could infect another seal which then travels for 13 days to reach Casey in mid-February. Another seal could be infected there and then take another 13 days to get to Commonwealth Bay arriving in late-February. A final seal in the chain could then be infected at Commonwealth Bay and take 15 days to reach Macquarie arriving in mid-March. This transmission chain is theoretically possible based on seal travel speeds and fits with the haulout schedules of both populations.

Little is known about when the virus has actually arrived at the other colonies, but elephant seals are known to be ashore all year, (Hindell & Burton 1988; McMahon, Burton & Bester 1999) so the virus could be locally maintained year-round. However other problems with the terrestrial stepping stone scenario are: (1) The transport host seals must be infectious on arrival but maintain high rates of travel on each step despite infection, (2) Commonwealth Bay is not a common haulout, (3) this chain requires 6 steps (or 5 if we begin from Heard Island), each of which is relatively unlikely, given the small number of seals that use the Antarctic mainland haulouts. The accumulated likelihood of 6 already low probability events suggests that this a possible but unlikely route. This is however tempered somewhat by the large population size at Kerguelen.

A series of terrestrial steps could not, however provide a plausible direct seal-to-seal transmission route to explain the observed spread from South Georgia to Kerguelen. There are no known continental haulouts between South Georgia and Marion Is, so the only intermediate step would be Bouvet Is, 2,560 km, requiring an infectious period of 19 days. The step from Bouvet to Marion Island is a further 2,536 km requiring an infectious period of 18 days. These large distances make seal-to seal transmission via terrestrial intermediaries unlikely between South Georgia and Kerguelen. However, there are feasible terrestrial stepping stones of the virus to move from Macquarie Is to New Zealand via seal-to-seal transmission.

### Possible at-sea routes of transmission

The observed dataset where two seals were at sea during the same 6-hour time step contained > 5,000,000 possible pairwise combinations. Restricting this to instances where pairs of seals were possibly in the same place at the same time (*i.e.* where their estimated location error ellipses overlapped) identified 33,285 possible at-sea encounters and these tended to be concentrated on the Kerguelen Plateau and on the Antarctic Continental Shelf, East of Prydz Bay (Figure 3a). Only one of these possible encounters was between seals from different breeding colonies. In this case, tracked seals from Kerguelen and Macquarie were in the same place at the same time (6h window) at a location off the coast from Commonwealth Bay (Figure3a).

In contrast, the population level analysis produced > 100,000,000 temporally congruent pairwise combinations, containing 200,661 potential at-sea encounters, 116 of which were between seals from different colonies (Figure 3b). The intra-colony encounters were widespread within the overall study domain, but were concentrated on the Kerguelen Plateau, the Antarctic continental shelf and north of the Ross Sea. The inter-colony encounters were primarily along the East Antarctic shelf (Macquarie/Kerguelen) or the western Weddell Sea (Marion/Kerguelen). This demonstrates that there is some potential for the virus to move from seal to seal during at-sea encounters, possibly via a series of both intra- and inter-colony interactions.

To better characterise the nature of these at-sea encounters we quantified how often each seal encountered another seal while at sea (Figure 4a). The frequency distribution was left-skewed with most seals encountering fewer than 5 other seals during their time at sea, but a small number of individuals encountered 20 or more other seals, demonstrating the potential for seals with high network connectivity (high contact rates with other animals) to act as “super-spreaders” within the population. The number of encounters between a pair of seals (a proxy for the duration and/or closeness of the encounter) showed a similar distribution (Figure 4b), with the majority of seals being in contact for less than 4 time steps (one day), but in some circumstances the seals were together for more than 12 time steps (3 days), which would provide increased likelihood of the type of close contact that would permit viral transmission, occurring between those two seals. The time between encounters (a measure of the likelihood of a seal encountering another while still infectious), showed that most of the encounters were less than 14 days apart (Figure 4c) - a time duration during which infected seals could potentially pass on the virus. Similarly, the majority of the encounters were less than 500 km apart (Figure 4d), suggesting a high likelihood of individuals remaining infectious from one encounter to the next.

**Figure 4:**
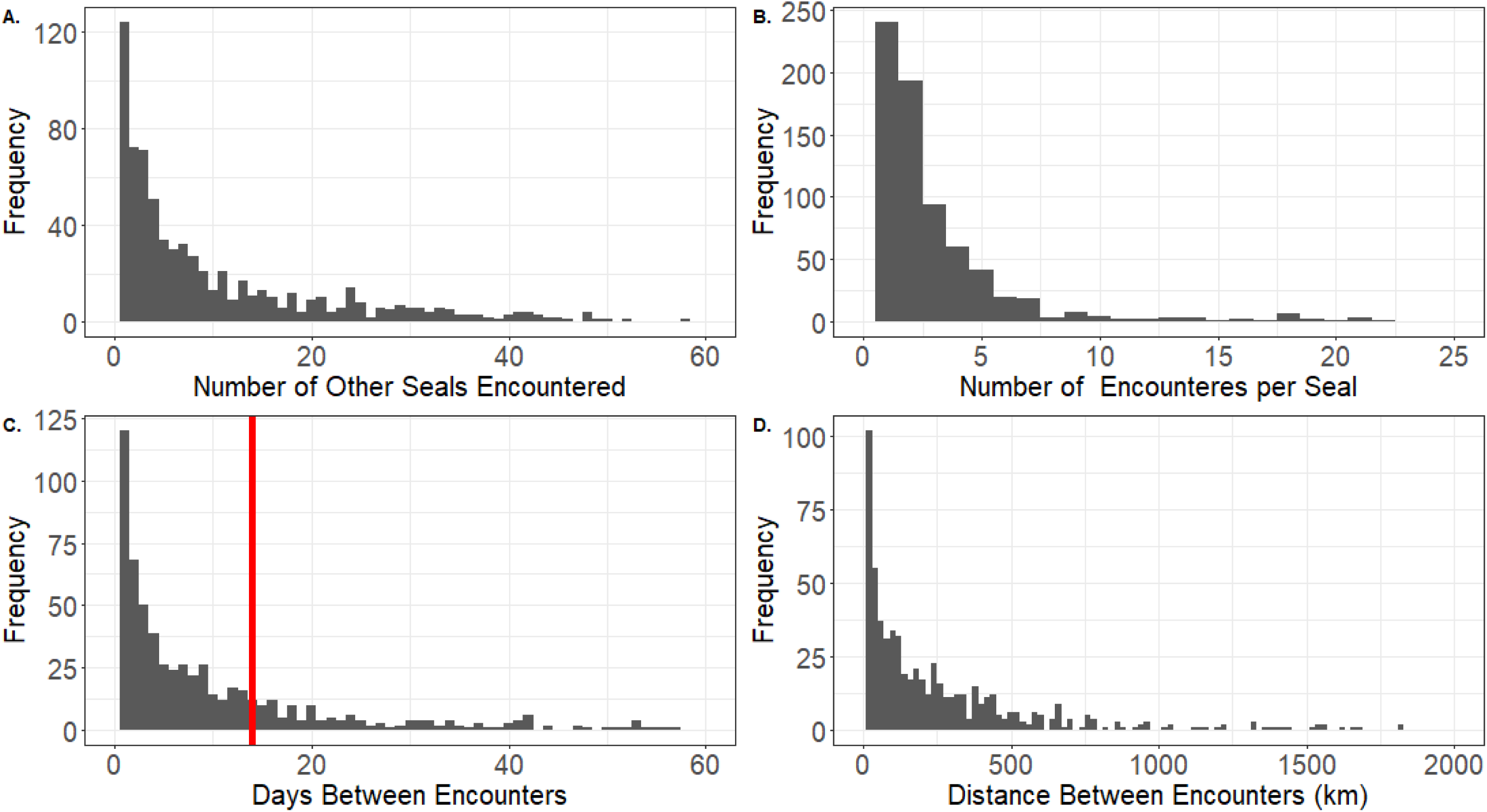
Descriptive statistics for the results of the seal at-sea encounters generated by the population level analysis. The figures present frequency distributions of summaries of 4 parameters for each of the 681 seals that had at least one at-sea encounter with another seal. The analyses were restricted to only encounters more than 200km from the breeding colony to avoid the large number of encounters that inevitably occur near the colonies. A. The number of other seals that each seal encountered at sea. B. The number of encounters with each of those seals, C. The mean time between successive encounters, where the red line is at 14 days indicating the period of plausible transmission and D. The mean distance (km) between each successive encounter.

From the tracking data, we identified two examples of how a virus might be transmitted between two breeding colonies via at-sea transmission (Figure 5). In both cases the seals had numerous at-sea encounters with other seals from their own colony before encountering a seal from a different colony, suggesting a possible transmission route between the colonies via at-sea encounters.

**Figure 5.**
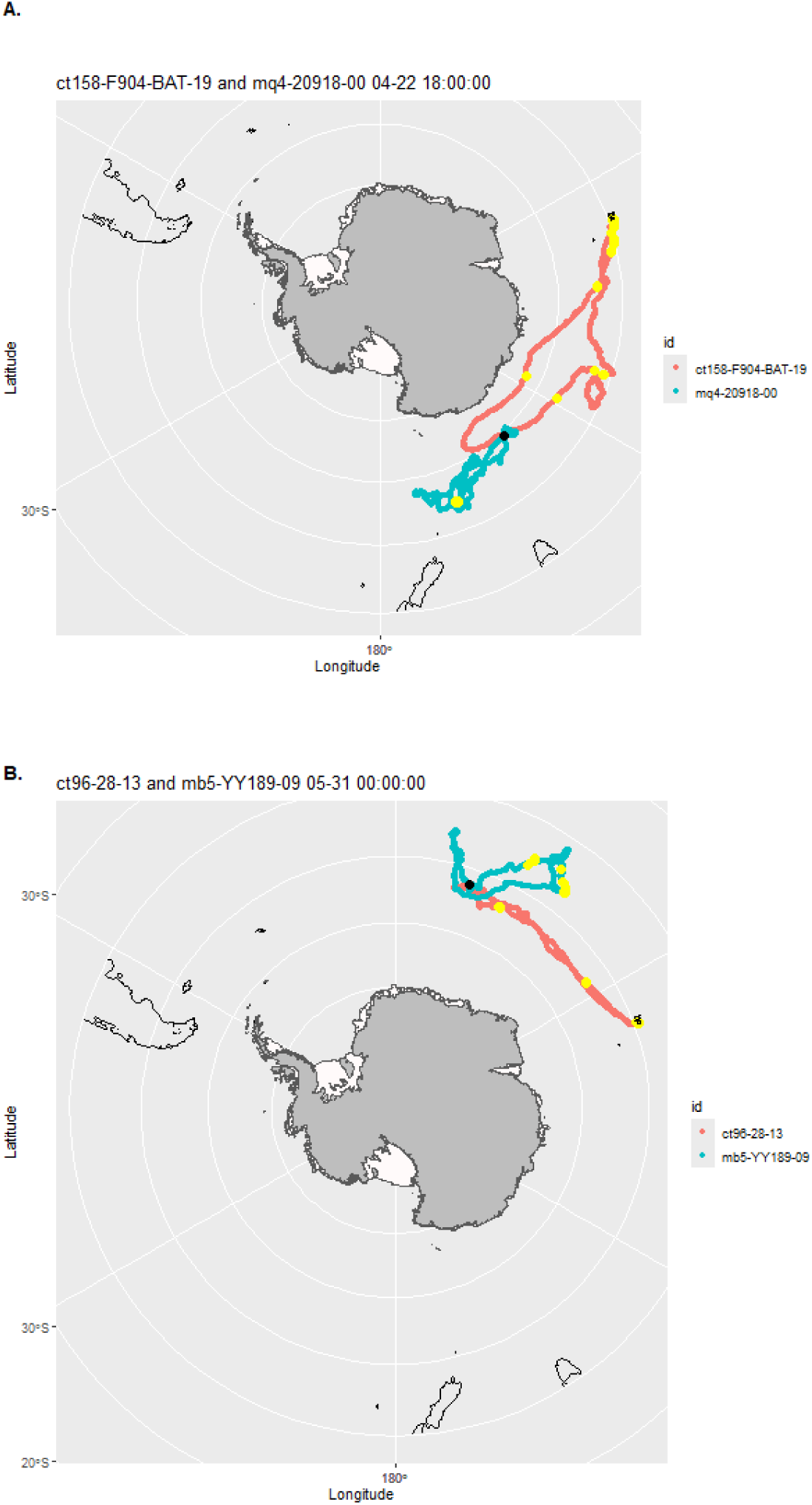
Two examples of tracks from the population level analysis where two seals from different breeding colonies potentially had at-sea encounters. The black dots indicate the location of the potential encounters between seals from different populations and the yellow dots indicate other encounters with seals from the same colony. In both cases the seals had numerous at-sea encounters with other seals from their own colony suggesting a possible transmission route between the colonies. A. seals ct158-F904-BAT-19 (From Iles Kerguelen) and mq4_20918_00 (from Macquarie Island) at 06:00 on 14 April, B. seals ct96_28_19 (Iles Kerguelen) and mb5_YY189_09 on May 31 at 00:00:00.

## Discussion

Seals, along with the broader Southern Ocean faunal communities, participate in tightly linked food webs and infectious disease can affect survival, breeding success and foraging behaviour in ways that can ripple through the ecosystem. Given the important role elephant seals play as major consumers in the Southern Ocean (Boyd, Arnbom & Fedak 1994; McMahon *et al*. 2019; Hindell *et al*. 2022), understanding how HPAI H5N1 affects their populations is central to better understanding the broader Southern Ocean ecosystem. The role seals may play in the spread of infectious agents like HPAI H5N1 require knowledge of incubation periods, shedding duration, length of the infectious period and the basis reproduction number, *R*_0_, all of which remain unknown for wild pinnipeds.

While polar ecosystems are characterized by their geographical isolation, which may constrain disease introduction, sustained mammal-to-mammal disease transmission (Xie *et al*. 2023) and spillover events remain plausible, particularly at coastal interfaces and densely packed colonies where seabirds and seals breed, moult and haul out (Hindell & Burton 1988; McMahon, Burton & Bester 1999). Using a large at-sea tracking data set from three elephant seal colonies in the Southern Indian and Pacific sectors of the Southern Ocean, we explored the feasibility of HPAI H5N1 reaching Macquarie Island via seal-to-seal transmission. We identified two transmission models, the “Terrestrial Stepping Stone Model”, and the “At-sea Transmission Model” (Figure 6). While both are possible when considering known rates of travel of elephant seals and likely infectious periods (as documented in other species), we conclude that the terrestrial stepping stone model is less likely, because it requires a series of relatively uncommon events to occur. By contrast, the at-sea transmission model is more likely due to the existence of commonly used foraging grounds and the large number of seals within the population. Our description of potential terrestrial and at-sea seal-to-seal interactions helps to understand population connectivity and co-location of probability surfaces (*e.g.* overlapping kernel densities like those we presented) and the exposure risk landscape. Furthermore, the findings allow us to determine the temporal dynamics of the overlap, all of which are important in predicting disease entry, establishment and spread.

**Figure 6:**
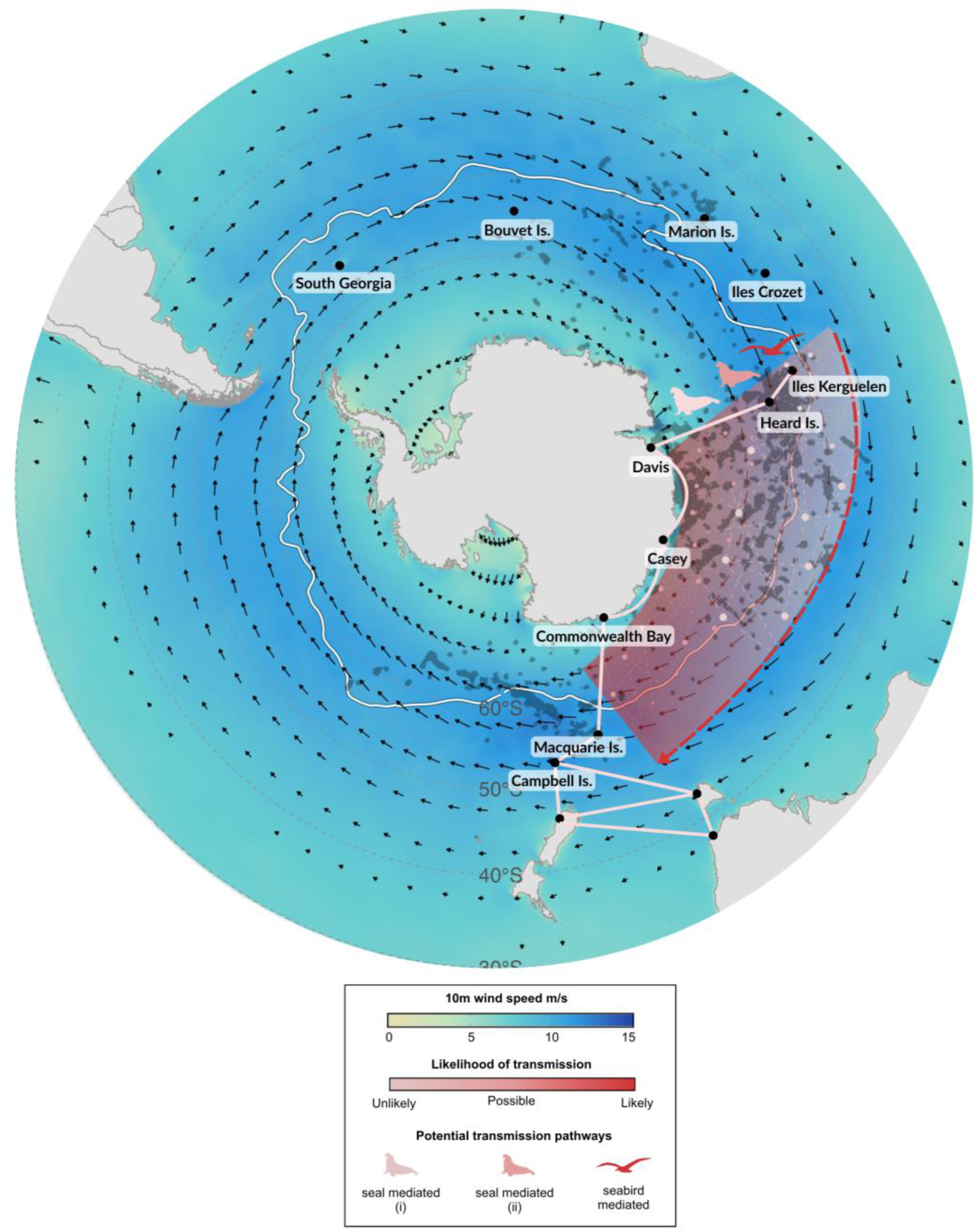
Schematic illustrating potential transmission pathways for HPAI from the southern Indian Ocean (*e.g.* Heard Island, the easternmost sub-Antarctic detection to date; June 2026) to the southwest Pacific Ocean, including Macquarie and Campbell Islands. Pathways are colour-coded by relative likelihood, with dark red indicating the highest likelihood and pale pink indicating lower likelihood. The study evaluated two seal-mediated transmission pathways: unlikely terrestrial “stepping-stone” movements between colonies (shown pale pink, limited to short routes), and more plausible at-sea transmission via inter-individual encounters (pink lattice, likelihood decreasing offshore). However, seabirds particularly circumpolar Procellariiformes and scavenging Charadriiformes are considered the most likely vectors overall, given long-distance movements, at-sea interactions, and scavenging behaviour, representing the highest-likelihood pathway (shown in dark red). The background shading represents mean ERA5 10 m wind speed from January 2024 to December 2025 (Copernicus Climate Change Service 2025), with arrows indicating the direction and relative magnitude of near-surface winds. These patterns reflect prevailing Southern Ocean circulation patterns from West to East, which may influence the movement and connectivity of potential biological vectors.

### HPAI H5N1 clade 2.3.4.4b longevity and time to move between haulout sites

Our models make two assumptions about HPAI H5N1 clade 2.3.4.4b transmission dynamics (i) a latent period of up to 4 days and (ii) an infectious period of up to 14 days. These values were chosen as biologically plausible working assumptions based on available influenza shedding data from humans, experimental mammal models, birds and cattle, while recognising that species-specific estimates are not currently available for pinnipeds. Experimental infections in several non-pinniped wild mammal and bird species show that nasal and oral shedding of HPAI H5N1 clade 2.3.4.4b can begin as early as one day post infection (Root *et al*. 2024). Infectious virus was detected through the first week in several mammal species, with the latest mammalian shedding detected at 8 days post-inoculation, while one bird remained positive at 14 days post-inoculation (Root *et al*. 2024). In cattle, viral load in milk samples following infection peaked rapidly, with minimum Ct values reached within 1-2 days, and the inferred duration of infectiousness had a central estimate of 6.2 days (95% credible interval: 2.8 to 13.1 days) (Eales, McCaw & Shearer 2025). Given that there are not marked differences in latent period or infectious period among birds, small mammals, cattle and humans, we expect these to be broadly similar in pinnipeds, indicating that H5N1 clade 2.3.4.4b infection in seals is likely to be characterised by a short latent period (no more than four days) and a fairly brief infectious window (no more than 14 days) for the vast majority of individuals. These parameters permit population-scale outbreaks to extend over weeks to months when ecological and behavioural conditions allow ongoing transmission across populations of susceptible host animals.

Field observations of HPAI H5N1-affected seal populations are consistent with a pattern of rapid disease progression at the individual level. Major mortality events in South America and the sub-Antarctic show that southern elephant seals and some seabird species typically succumb quickly after developing neurological and respiratory signs (Uhart *et al*. 2024; Beckmen *et al*. 2025; Knief *et al*. 2026). On South Georgia, Crozet and Kerguelen islands, large numbers of elephant seal pups were found dead or dying shortly after the onset of symptoms, with extremely high mortality rates recorded during the initial weeks following detection (Campagna *et al*. 2023; Clessin *et al*. 2025)^.(McInnes *et al*. 2026)^ At Península Valdés, mortality among southern elephant seal pups continued for at least six weeks, with peak mortality occurring between late September and mid-October and remaining elevated into November (Uhart *et al*. 2024; Beckmen *et al*. 2025; Ashley *et al*. 2026). Similar patterns were observed among harbour and grey seals in Maine USA, where peak levels of mortality lasted less than one month. By contrast outbreaks among South American sea lions have continued for more than eight months across the regional metapopulation (Leguia *et al*. 2023; Plaza *et al*. 2024; Puryear & Runstadler 2024; Tomas *et al*. 2024; Uhart *et al*. 2024). These latter prolonged outbreaks reflect sustained chains of transmission within and between colonies rather than protracted infectiousness in individual animals. While the occurrence of mass mortality events among marine mammals has been well documented, there remains much to be learnt. Reviews of mammal infections note that most marine mammals are only detected once they are severely ill or already dead, which limits scientific opportunities to characterise early infection dynamics, quantify the latent and infectious periods and determine prevalence estimates(Plaza *et al*. 2024). Detecting infection in free-ranging marine mammals is challenging, but at least a small number of asymptomatic mild infections are likely to occur. Active surveillance of apparently healthy animals is therefore essential to identify ongoing infections and virus shedding. Molecular testing for current infection and serological testing for previous exposure are crucial understanding transmission dynamics in susceptible species like pinnipeds and birds.

### The terrestrial transmission of HPAI H5N1 clade 2.3.4.4b

Oceania long remained free of HPAI H5N1 – up until June 2026. This may be due its isolation from regions where the virus has been detected because the distances that infected animals must cross to reach these land masses may be too great if infected animals die *en route* during these long migrations. Short-tailed shearwaters travelling between the Arctic and Australia, for example, cover approximately 11,000 km in under two weeks (Carey *et al*. 2014; Bool *et al*. 2024) a feat which is likely made much more difficult in the presence of viral infection. There are also relatively few such species making these long-distance migrations. While there is high connectivity from Australia (and to a lesser extent New Zealand) to eastern Indonesia and Papua New Guinea, there is limited connectivity to broader southeast Asia beyond Wallace’s Line (McCallum *et al*. 2008; Gilbert *et al*. 2010). The circulation of HPAI in the Antarctic and sub-Antarctic creates a new dynamic environment for incursion risk to HPAI-free countries in Oceania with different species involved.

While there exists a series of connected haulout sites between the major elephant seal populations, terrestrial spread seems unlikely, given the seals can only cover the distances between key haulout sites within the infectious period of up to 14 days via a complex and improbable series of exposure and transmission events. Animal tracking data represents only a subset of individuals, potentially biasing population estimates. Scaling fine-scale behavioural data to population-level epidemiological dynamics is not straightforward. We aim to estimate transmission pathway plausibility rather than precise probabilities.

Ecosystems contain multiple species and pathogens acting as hosts and reservoirs. Integrating information from across the diversity of species remains a frontier challenge given not all marine species are equally well studied (Hindell *et al*. 2020; Sequeira *et al*. 2025) and neither are pathogen communities, indeed in most cases these are almost completely unknown.

### At sea transmission of H5N1

At-sea transmission of HPAI H5N1 in elephant seals represents a complex yet poorly understood potential pathway of disease spread. Elephant seals undertake long pelagic migrations lasting up to eight months to forage across oceanic regions (Hindell *et al*. 2016), with extensive regions of overlap between populations. During their time at sea, seals are, as we show, at times near conspecifics, including those from other populations creating opportunities for interaction with contaminated surface waters, floating carcasses, or direct contact. Although influenza viruses tend to degrade more rapidly in saline than in freshwater environments, low ocean temperatures, reduced ultraviolet radiation at high latitudes in the winter, and organic material in the water column may enable viral persistence long enough to permit infection if contact is close in both space and time (Van Bressem *et al*. 2009; Brown *et al*. 2014; Keeler *et al*. 2014; Kaiser *et al*. 2025). Direct transmission may also occur if elephant seals investigate or scavenge infected carcasses, a behaviour that, while not typical, has been documented in some pinniped species under opportunistic feeding conditions (Robinson *et al*. 2012). Furthermore, elephant seals are not entirely solitary at sea; they may aggregate in areas of high prey density such as upwelling zones, seamounts, or frontal systems, where temporary clustering could facilitate respiratory or close-contact transmission between individuals (Hindell *et al*. 2020). Surface social interactions; vocalizations, physical contact, and communal foraging may facilitate aerosol/droplet transmission of H5N1 among seals. At-sea transmission could spread the virus to distant colonies without involvement of terrestrial haul-outs, complicating surveillance since infected individuals may show no symptoms until returning to land. Understanding at-sea contact patterns is crucial for predicting outbreak trajectories, population impacts, and cross-species transmission risks to other marine mammals or birds.

A likely point of contact for the seals at-sea would be on ice floes. Sea-ice is concentrated across the Antarctic continental shelf, extending out into the deeper ocean as the winter progresses. The ice provides a stable platform on which the seals can haul out to rest and may act as a catalyst for individuals aggregating and interacting. While there is no evidence that seals forage in groups or communally, our analysis suggests that some individuals encounter up to 20 other seals, and they may stay together for up to 3 days. If this were to happen in the sea-ice the potential for transmission would be high. To date, at-sea haulout behaviours remain poorly known, as is how seals encountering each other in open water would interact, but it is likely that the at-sea interactions are more transitory than when sharing an ice floe. Indeed, such interactions on ice floes provide a transmission route into other species of seal, crabeater (*Lobodon carcinophagus)*, leopard (*Hydrurga leptonyx)*, Weddell (*Leptonychotes weddellii*) and Ross (*Ommatophoca rossii*) seals that also use the ice as haulout and resting platforms (Younger *et al*. 2026).

We used a simple analysis to test the At-sea Transmission model, by assuming all 791 tag deployments were made in the same year, rather than spread over a 20-year period. In so doing we are still likely to greatly underestimate the rate of at-sea encounters, as our sample of nearly 800 seals is only a small fraction of the total number of seals in the populations (350,000 for Iles Kerguelen (Laborie *et al*. 2023), 40,110 at Macquarie Island and 5,500 for Marion Island (Oosthuizen *et al*. 2015; Laborie *et al*. 2023; Hindell *et al*. 2025)). Consequently, at sea encounters are likely to be even more common than we describe and offer an alternative route of transmission between sub-Antarctic colonies of elephant seals. This suggests two related pathways for the virus to move seal-to-seal. First, the virus could move between colonies via a series of at-sea encounters (see Figure 6). Alternatively, the at-sea encounters might maintain a pool of infected seals away from the colonies throughout the winter months, a source from which infected animals arrive at new islands in the breeding season. Full examination of these (and other) alternatives will require more detailed epidemiological models than are possible in this exploratory study. Nevertheless, we have demonstrated that seal-to-seal transmission is a plausible mechanism for viruses such as HPAI H5N1 to move between even remote sub-Antarctic Islands and emphasise the potential value of integration of long term population monitoring and movement data with understanding disease dynamics in wildlife. Our demonstration of the frequency of at-sea encounters also suggests that there is a possibility of at-sea mortality which is currently unaccounted for in population level assessments of disease effects.

### Seabird-mediated transmission

Although this study describes plausible pathways of the spread of HPAI H5N1 by southern elephant seals in Antarctica and the Southern Ocean, the most likely pathway for introduction, establishment and spread west into Australasia remains via wide-ranging seabirds (Figure 7). Based on the movement patterns of seabirds along the Southern Ocean Flyway (Morten *et al*. 2025)^,^ their central position suggests Macquarie Island and New Zealand are likely to eventually be visited by potential seabird hosts with documented connectivity to known HPAI H5N1 affected regions of the Southern Ocean (Williams *et al*. 2006; Leguia *et al*. 2023; Xie *et al*. 2023; Wille *et al*. 2024a). Among these, the southern giant petrel poses perhaps the greatest incursion risk while other species such as the Brown Skua, white-headed petrel (Stanislawek *et al*. 2024; Wille *et al*. 2024a), Campbell albatross (Thompson *et al*. 2021) and grey petrel (Torres *et al*. 2015) may also contribute to the risk. Both HPAI positive giant petrels and skuas were recently detected in Australia, and are also likely to reach New Zealand or Macquarie Island which supports more than three million seabirds from over 57 species (Cresswell *et al*. 2023). As with the seal-mediated pathway we discussed here, long-distance transmission by seabirds also requires a confluence of conditions including infection durations that exceed the transit time and thereby support viral shedding upon arrival. The ongoing incursion risk to Macquarie Island and New Zealand has been assessed as low (Van Bressem *et al*. 2009; Fountain-Jones *et al*. 2024; Wille *et al*. 2024a; Wille *et al*. 2024b), although the recent detections in Australia show that this is developing rapidly. The ecological connectivity demonstrated here underscores the need for continued vigilance as both the panzootic and the virus evolve.

## Conclusion

Elephant seals could transmit H5N1 between sub-Antarctic islands during their 8-month marine phase if infected individuals remain mobile and infectious without showing clinical signs. Tracking data from three colonies indicates this viral transmission pathway is biologically plausible based on encounter rates and travel speeds, although the key uncertainty involves whether infected seals maintain mobility despite infection (Sobolev *et al*. 2024; Alekseev *et al*. 2025). Since the existence (or non-existence) of subclinical or mild cases in elephant seals is central to this pathway, active sampling of apparently healthy animals, including molecular testing for current infection and serology for previous exposure is warranted. Long-distance dispersal by birds remains the most likely pathway, but seal-mediated transmission should not be excluded without better data on infection outcomes, shedding duration and survival in free-ranging pinnipeds. Resolving the relative contribution of avian and mammalian hosts will require integrated movement data, active surveillance of apparently healthy animals, and thorough sampling to permit genomic analyses capable of distinguishing new introductions from local maintenance (Sobolev *et al*. 2024; Alekseev *et al*. 2025).

The combination of ecological vulnerability (dense colonial breeding aggregations), remoteness (causing delayed detection and responses), multiple transmission pathways and viral evolution mean these outbreaks are likely to continue and the impacts could substantially reshape sub-Antarctic and Antarctic marine ecosystems. Surveillance, cross-disciplinary research using a One Health approach (Fountain-Jones *et al*. 2024), and stringent biosecurity controls to prevent human-induced spread are pragmatic actions that help understand and mitigate impacts. Given the difficulty of implementing broad-scale disease control in remote wildlife populations, direct interventions to reduce HPAI H5N1 mortality may only be feasible in narrowly targeted contexts, such as vaccination of high-risk species in small populations. Broader conservation efforts should therefore focus on supporting the resilience and recovery of impacted or at-risk wildlife populations. Developing an integrated epidemiological model for the Southern Ocean and Antarctica, including consideration of the role of communal sea-ice seal haulouts and sex-specific behaviour in transmission pathways would be valuable, as would on-going monitoring of the molecular evolution and epidemiology of the virus.

The surveillance, understanding and management of wildlife diseases remain challenging (Woods *et al*. 2019). This study demonstrates some of the difficulties of assessing risk, transmission and the likelihood of entry, establishment and spread of viruses in wild animal populations and contributes to a better understanding of the epidemiology of HPAI H5N1 in the Southern Ocean. Moreover, our study demonstrates the value of integrating long-term population monitoring and movement data into our understanding of infectious disease epidemiology in wildlife which can be especially useful for understanding epidemiology of high-consequence viruses (Rupprecht *et al*. 2018; Rupprecht & Belsare 2026; Gridley *et al*. PREPRINT).

## Acknowledgements

The seal tracking study was supported by the Integrated Marine Observing System (IMOS) – IMOS is enabled by the National Collaborative Research Infrastructure Strategy (NCRIS) and the IPEV programs no. 109 (PI H. WEIMERSKIRCH) and no. 1201 (PI C. GILBERT). Fieldwork at the Australian Antarctic stations was supported by the Australian Antarctic Division (AAS 2265, AAS 2794, AAS 4329, AAS 4605, & AAS 4630). The Australian Research Council supported this work through Discovery Projects: DP0345010, DP0770910, DP180101667, & DP23010136 and the ARC Special Research Initiative SR200100008. All tagging procedures were approved and executed under University of Tasmania Animal Ethics Committee guidelines (A12141, A14523), the Comité d’éthique Anses/ENVA/ UPEC (no. APAFiS: 21375), and by Macquarie University Ethics Committee ARA 2014_057. We thank Drs. Marcela Uhart and Andrew Breed for their thoughtful and helpful reviews.

## References

Alekseev, A., Sobolev, I., Sharshov, K., Gulyaeva, M., Kurskaya, O., Kasianov, N., Chistyaeva, M., Ivanov, A., Ohlopkova, O., Moshkin, A., Stepanyuk, M., Derko, A., Solomatina, M., Mutashev, B., Dolgopolova, M., Gadzhiev, A. & Shestopalov, A.; 2025; Pathobiology of Highly Pathogenic Avian Influenza A (H5N1 Clade 2.3.4.4b) Virus from Pinnipeds on Tyuleniy Island in the Sea of Okhotsk, Russia; Viruses; 1810.3390/v18010051.

Argos; 2016; Argos User’s Manual: Worldwide Tracking and Environmental Monitoring by Satellite; Toulouse: Argos.

Ashley, E., Vanstreels, R.E.T., Barbieri, M., Puryear, W., Gulland, F., Field, C., Johnson, C.K. & Uhart, M.; 2026; High pathogenicity avian influenza in pinniped conservation; Philosophical Transactions of the Royal Society B: Biological Sciences 38110.1098/rstb.2024.0320.

Bamford, C.C.G., Fenney, N., Coleman, J., Fox-Clarke, C., Dickens, J., Fedak, M., Fretwell, P., Hückstädt, L. & Hollyman, P.; 2025; Highly Pathogenic Avian Influenza Viruses (HPAIV) Associated with Major Southern Elephant Seal Decline at South Georgia; Communications Biology; 810.1038/s42003-025-09014-7.

Beckmen, K.B., Burek Huntington, K.A., Spraker, T. & Ellis, J.; 2025; Pathologic Characterization of Highly Pathogenic H5N1 Avian Influenza Virus Infections in Wild Mammals in Alaska, USA; Journal of Wildlife Disease; 10.7589/JWD-D-25-00023.

Bester, M.N.; 1988; Marking and monitoring studies of the Kerguélen stock of southern elephant seals *Mirounga leonina* and their bearing on biological research in the Vestfold Hills; Hydrobiologia; 165; 269–277;

Bester, M.N., Bornemann, H., Daneri, G.A. & van den Hoff, J. (2020) Southern elephant seals (*Mirounga leonina* L.) in the Antarctic Treaty Area. SCAR.

Bi, Y., Yang, J., Wang, L., Ran, L. & Gao, G.F.; 2024; Ecology and evolution of avian influenza viruses; Current Biology; 34; R716–R721; 10.1016/j.cub.2024.05.053.

Bool, N., Sumner, M.D., Lea, M.-A., McMahon, C.R. & Hindell, M.A.; 2024; Hierarchical foraging strategies of migratory short-tailed shearwaters during the non-breeding stage; Marine Biology; 17110.1007/s00227-023-04370-6.

Boyd, I.L., Arnbom, T. & Fedak, M.A. (1994) Biomass and energy consumption of the South Georgia population of southern elephant seals. Elephant seals: population ecology, behaviour, and physiology (eds B.J. Le Boeuf & R.M. Laws), pp. 98–117. University of California Press, Berkeley.

Brown, J., Stallknecht, D., Lebarbenchon, C. & Swayne, D.; 2014; Survivability of Eurasian H5N1 highly pathogenic avian influenza viruses in water varies between strains; Avian Diseases; 58; 453–457; 10.1637/10741-120513-ResNote.1.

Campagna, C., Uhart, M., Falabella, V., Campagna, J., Zavattieri, V., Vanstreels, R.E.T. & Lewis, M.N.; 2023; Catastrophic mortality of southern elephant seals caused by H5N1 avian influenza; Marine Mammal Science; 10.1111/mms.13101.

Carey, M.J., Phillips, R.A., Silk, J.R.D. & Shaffer, S.A.; 2014; Trans-equatorial migration of Short-tailed Shearwaters revealed by geolocators; Emu; 114; 352–359; 10.1071/MU13115.

Chua, M., Ho, S.Y.W., McMahon, C.R., Jonsen, I.D. & de Bruyn, M.; 2022; Movements of southern elephant seals (Mirounga leonina) from Davis Base, Antarctica: combining population genetics and tracking data; Polar Biology; 45; 1163–1174; 10.1007/s00300-022-03058-9.

Clessin, A., Briand, F.-X., Tornos, J., Lejeune, M., De Pasquale, C., Fischer, R., Souchaud, F., Hirchaud, E., Hong, S.L., Bralet, T., Guinet, C., McMahon, C.R., Grasland, B., Baele, G. & Boulinier, T.; 2025; Circumpolar spread of avian influenza H5N1 to southern Indian Ocean islands; Nature Communications; 1610.1038/s41467-025-64297-y.

Clessin, A., Brusselmans, M., Hong, S.L., Tornos, J., Lejeune, M., Shao, Y., Briand, F.X., Abolnik, C., Kaza, B., Suchard, M.A., Aguado, B., Alcami, A., Barbraud, C., Beer, M., Bennison, A., Bonadonna, F., Bonnet, T., Bost, C.A., Boucheron, S., Bralet, T., Catry, P., Cleeland, J., Connan, M., Coombes, H.A., Delord, K., de Pasquale, C., Dewar, M.L., Dong, X., Emerit, J., Fischer, R., Fountain-Jones, N., Galimberti, F., Gonzalez-Solis, J., Guinet, C., Gunn, C., Gunther, A., Iervolino, M., James, J., Ji, X., Jonsen, I., Jones, C.W., Kuepfer, A., Kuiken, T., Lebohec, C., Lisovski, S., Llorente Zubiri, L., Lynton-Jenkins, J.G., Martinez-Garcia, P., McCulley, M., McMahon, C.R., Mollett, B.C., Moraga-Quintanilla, A.I., Nichol, R., Noiret, A., Ogrzewalska, M., Owen, K., Pardo-Roa, C., Peroteau, S., Phillips, R.A., Poulin, E., Rambaut, A., Reid, S.M., Riehle, E., Risi, M., Rumianowski, O., Ryan, P.G., Sanvito, S., Stanworth, A., Steinfurth, A., Stevenson, J., Stier, A., Uhart, M.M., Vanstreels, R.E.T., Vazquez-Calvo, A., Vianna, J.A., Wells, K., White, J., Whitelaw, P., Wille, M., Younger, J., Roberts, L.C., Grasland, B., Banyard, A.C., Nelson, M.I., Gamble, A., Boulinier, T. & Baele, G.; 2026; Dispersal, adaptation and persistence of H5N1 in the sub-Antarctic and Antarctica; bioRxiv; 10.64898/2026.03.20.713283.

Copernicus Climate Change Service; 2025; ERA5 hourly time-series data on single levels from 1940 to present; Copernicus Climate Change Service (C3S) Climate Data Store (CDS); 10.24381/cds.adbb2d47

Cresswell, I., Bax, N., Constable, A., Reid, K. & Smith, A.M. (2023) The unique marine ecosystem surrounding Macquarie Island. Australian Marine Conservation Society.

de Bruyn, P.J.N., Tosh, C.A., Bester, M.N., Cameron, E.Z., McIntyre, T. & Wilkinson, I.S.; 2011; Sex at sea: alternative mating system in an extremely polygynous mammal; Animal Behaviour; 82; 445–451; 10.1016/j.anbehav.2011.06.006.

Eales, O., McCaw, J.M. & Shearer, F.M.; 2025; Viral kinetics of H5N1 infections in dairy cattle; 10.1101/2025.02.01.636082.

Field, I.C., Bradshaw, C.J.A., McMahon, C.R., Harrington, J. & Burton, H.R.; 2002; Effects of age, size and condition of elephant seals (*Mirounga leonina*) on their intravenous anaesthesia with tiletamine and zolazepam; Veterinary Record; 151; 235–240;

Field, I.C., Harcourt, R.G., Boehme, L., de Bruyn, P.J.N., Charrassin, J.B., McMahon, C.R., Bester, M.N., Fedak, M.A. & Hindell, M.A.; 2012; Refining instrument attachment on phocid seals; Marine Mammal Science; 28; E325–E332; 10.1111/j.1748-7692.2011.00519.x.

Fielding, J.E., Kelly, H.A., Mercer, G.N. & Glass, K.; 2014; Systematic review of influenza A(H1N1)pdm09 virus shedding: duration is affected by severity, but not age; Influenza and other Respiritory Viruses; 8; 142–150; 10.1111/irv.12216.

Fountain-Jones, N.M., Hutson, K.S., Jones, M., Nowak, B.F., Turnbull, A., Younger, J., O’Reilly, M., Watkins, E., Guernier-Cambert, V., Cooley, L. & Hamede, R.; 2024; One Health on islands: Tractable ecosystems to explore the nexus between human, animal, terrestrial, and marine health; BioScience; 10.1093/biosci/biae101.

Gilbert, M., Newman, S.H., Takekawa, J.Y., Loth, L., Biradar, C., Prosser, D.J., Balachandran, S., Subba Rao, M.V., Mundkur, T., Yan, B., Xing, Z., Hou, Y., Batbayar, N., Natsagdorj, T., Hogerwerf, L., Slingenbergh, J. & Xiao, X.; 2010; Flying over an infected landscape: distribution of highly pathogenic avian influenza H5N1 risk in South Asia and satellite tracking of wild waterfowl; Ecohealth; 7; 448–458; 10.1007/s10393-010-0672-8.

Glynn, J.R., Bower, H., Johnson, S., Houlihan, C.F., Montesano, C., Scott, J.T., Semple, M.G., Bangura, M.S., Kamara, A.J., Kamara, O., Mansaray, S.H., Sesay, D., Turay, C., Dicks, S., Wadoum, R.E.G., Colizzi, V., Checchi, F., Samuel, D. & Tedder, R.S.; 2017; Asymptomatic infection and unrecognised Ebola virus disease in Ebola-affected households in Sierra Leone: a cross-sectional study using a new non-invasive assay for antibodies to Ebola virus; The Lancet Infectious Diseases; 17; 645–653; 10.1016/S1473-3099(17)30111-1.

Gridley, T., Barnes, A., Blumberg, L., Creighton, A., Dines, S., Donovan, A., Elwen, S., Frainer, G., Friedman, J., Gardner, B., Hofmeyr, G., Jenkinson, I., Kieswetter, N., Mendes, L., Nadin, C., Oelofse, G., Pieterse, J., Probert, R., Seakamela, S., Simpson, G., Steyl, J., Farley, E. & Kotze, J.; PREPRINT; Novel rabies outbreak in a marine mammal - the Cape fur seal,; [10.21203/rs.3.rs-7992979/v1]

Hindell, M.A. & Burton, H.R.; 1988; Seasonal haul-out patterns of the southern elephant seal (*Mirounga leonina*) at Macquarie Island; Journal of Mammalogy; 69; 81–88; 10.2307/1381750.

Hindell, M.A. & Little, G.J.; 1988; Longevity, fertility and philopatry of two female southern elephant seals (*Mirounga leonina*) at Macquarie Island; Marine Mammal Science; 4; 168–171; 10.1111/j.1748-7692.1988.tb00197.x.

Hindell, M.A. & McMahon, C.R.; 2000; Long distance movement of a southern elephant seal (*Mirounga leonina*) from Macquarie Island to Peter 1 Ø y; Marine Mammal Science; 16; 504–507;

Hindell, M.A., McMahon, C.R., Bester, M.N., Boehme, L., Costa, D., Fedak, M.A., Guinet, C., Herraiz-Borreguero, L., Harcourt, R.G., Huckstadt, L., Kovacs, K., M., Lydersen, C., McIntyre, T., Muelbert, M.M.C., Patterson, T., Roquet, F., Williams, G. & Charrassin, J.B.; 2016; Circumpolar habitat use in the southern elephant seal: implications for foraging success and population trajectories; Ecosphere; 7; e01213; 10.1002/ecs2.1213.

Hindell, M.A., McMahon, C.R., Guinet, C., Harcourt, R., Jonsen, I.D., Raymond, B. & Maschette, D.; 2022; Assessing the potential for resource competition between the Kerguelen Plateau fisheries and southern elephant seals; Frontiers in Marine Science; 910.3389/fmars.2022.1006120.

Hindell, M.A., McMahon, C.R., Harcourt, R., Arce, F., Guinet, C. & Jonsen, I.; 2021; Inter- and intra-sex habitat partitioning in the highly dimorphic southern elephant seal; Ecology and Evolution; 10.1002/ece1003.7147; 10.1002/ece3.7147.

Hindell, M.A., McMahon, C.R., Van Den Hoff, J., Thalmann, S., Carlyon, K. & Wotherspoon, S.; 2025; The ongoing decrease in numbers of breeding female southern elephant seals (Mirounga leonina L.) at Macquarie Island; Antarctic Science; 1–9; 10.1017/s0954102025000161.

Hindell, M.A., Reisinger, R.R., Ropert-Coudert, Y., Huckstadt, L.A., Trathan, P.N., Bornemann, H., Charrassin, J.B., Chown, S.L., Costa, D.P., Danis, B., Lea, M.A., Thompson, D., Torres, L.G., Van de Putte, A.P., Alderman, R., Andrews-Goff, V., Arthur, B., Ballard, G., Bengtson, J., Bester, M.N., Blix, A.S., Boehme, L., Bost, C.A., Boveng, P., Cleeland, J., Constantine, R., Corney, S., Crawford, R.J.M., Dalla Rosa, L., de Bruyn, P.J.N., Delord, K., Descamps, S., Double, M., Emmerson, L., Fedak, M., Friedlaender, A., Gales, N., Goebel, M.E., Goetz, K.T., Guinet, C., Goldsworthy, S.D., Harcourt, R., Hinke, J.T., Jerosch, K., Kato, A., Kerry, K.R., Kirkwood, R., Kooyman, G.L., Kovacs, K.M., Lawton, K., Lowther, A.D., Lydersen, C., Lyver, P.O., Makhado, A.B., Marquez, M.E.I., McDonald, B.I., McMahon, C.R., Muelbert, M., Nachtsheim, D., Nicholls, K.W., Nordoy, E.S., Olmastroni, S., Phillips, R.A., Pistorius, P., Plotz, J., Putz, K., Ratcliffe, N., Ryan, P.G., Santos, M., Southwell, C., Staniland, I., Takahashi, A., Tarroux, A., Trivelpiece, W., Wakefield, E., Weimerskirch, H., Wienecke, B., Xavier, J.C., Wotherspoon, S., Jonsen, I.D. & Raymond, B.; 2020; Tracking of marine predators to protect Southern Ocean ecosystems; Nature; 580; 87–92; 10.1038/s41586-020-2126-y.

Jonsen, I.D., Grecian, W.J., Phillips, L., Carroll, G., McMahon, C., Harcourt, R.G., Hindell, M.A. & Patterson, T.A.; 2023; aniMotum, an R package for animal movement data: Rapid quality control, behavioural estimation and simulation; Methods in Ecology and Evolution; 10.1111/2041-210x.14060.

Kaiser, F., Cardenas, S., Yinda, K.C., Mukesh, R.K., Ochwoto, M., Gallogly, S., Wickenhagen, A., Bibby, K., de Wit, E., Morris, D., Lloyd-Smith, J.O. & Munster, V.J.; 2025; Highly Pathogenic Avian Influenza A(H5N1) Virus Stability in Irradiated Raw Milk and Wastewater and on Surfaces, United States; Emerging Ingfectious Diseases; 31; 833–837; 10.3201/eid3104.241615.

Keeler, S.P., Dalton, M.S., Cressler, A.M., Berghaus, R.D. & Stallknecht, D.E.; 2014; Abiotic factors affecting the persistence of avian influenza virus in surface waters of waterfowl habitats; Applied and Environmental Microbiolgy; 80; 2910–2917; 10.1128/AEM.03790-13.

Knief, U., Bouwhuis, S., Globig, A., Gunther, A. & Courtens, W.; 2026; Counting cases, conserving species: addressing highly pathogenic avian influenza in wildlife; Biological reviews; 10.1002/brv.70173.

Kuiken, T., Vanstreels, R.E.T., Banyard, A., Begeman, L., Breed, A.C., Dewar, M., Fijn, R., Serafini, P.P., Uhart, M. & Wille, M.; 2026; Emergence, spread, and impact of high-pathogenicity avian influenza H5 in wild birds and mammals of South America and Antarctica; Conservtion Biology; 40; e70052; 10.1111/cobi.70052.

Laborie, J., Authier, M., Chaigne, A., Delord, K., Weimerskirch, H. & Guinet, C.; 2023; Estimation of total population size of southern elephant seals (Mirounga leonina) on Kerguelen and Crozet Archipelagos using very high-resolution satellite imagery; Frontiers in Marine Science; 1010.3389/fmars.2023.1149100.

Leguia, M., Garcia-Glaessner, A., Munoz-Saavedra, B., Juarez, D., Barrera, P., Calvo-Mac, C., Jara, J., Silva, W., Ploog, K., Amaro, L., Colchao-Claux, P., Johnson, C.K., Uhart, M.M., Nelson, M.I. & Lescano, J.; 2023; Highly pathogenic avian influenza A (H5N1) in marine mammals and seabirds in Peru; Nature Communications; 14; 5489; 10.1038/s41467-023-41182-0.

McCallum, H.I., Roshier, D.A., Tracey, J.P., Joseph, L. & Heinsohn., R.; 2008; Will Wallace’s Line save Australia from avian influenza? 13(2): 41; Ecology and Society 13; Article 41;

McInnes, J.C., Burgess, T., Mergard, G., Wells, M.R., McMahon, C.R., Neave, M.J., Polanowski, A., Terauds, A., Tornos, J., Lejeune, M., Briand, F.-X., Baele, G., Boulinier, T., Achurch, H., Alderman, R., Lashko, A., Wienecke, B., Wynen, L.P., Viola, B., Virtue, P. & Hodgson, J.C.; 2026; Mass mortality of southern elephant seals during multi-species outbreak of HPAI H5N1 on sub-Antarctic Heard Island; 10.64898/2026.06.16.732752.

McMahon, C.R., Burton, H.R. & Bester, M.N.; 1999; First-year survival of southern elephant seals, *Mirounga leonina*, at sub-Antarctic Macquarie Island; Polar Biology; 21; 279–284;

McMahon, C.R., Burton, H.R., McLean, S., Slip, D. & Bester, M.N.; 2000; Field immobilisation of southern elephant seals with intravenous tiletamine and zolazepam; Veterinary Record; 146; 251–254;

McMahon, C.R., Field, I.C., Bradshaw, C.J.A., White, G.C. & Hindell, M.A.; 2008; Tracking and data-logging devices attached to elephant seals do not affect individual mass gain or survival Journal of Experimental Marine Biology and Ecology; 360; 71–77; 10.1016/j.jembe.2008.03.012.

McMahon, C.R., Hindell, M.A., Charrassin, J.B., Corney, S., Guinet, C., Harcourt, R., Jonsen, I., Trebilco, R., Williams, G. & Bestley, S.; 2019; Finding mesopelagic prey in a changing Southern Ocean; Scientific Reports; 9; 19013; 10.1038/s41598-019-55152-4.

McMahon, C.R., Roquet, F., Guinet, C., Hindell, M.A., Harcourt, R., Charrassin, J.-B., Labrousse, S., Jonsen, I., Picard, B., Bestley, S., Doriot, V. & Fedak, M.; 2025; An enduring, 20-year, multidisciplinary seal-borne ocean sensor research collaboration in the Southern Ocean; Elementa Science of the Anthropocene; 13; 00071; 10.1525/elementa.2024.00071.

Morten, J.M., Carneiro, A.P.B., Beal, M., Bonnet-Lebrun, A.S., Dias, M.P., Rouyer, M.M., Harrison, A.L., González-Solís, J., Jones, V.R., Garcia Alonso, V.A., Antolos, M., Arata, J.A., Barbraud, C., Bell, E.A., Bell, M., Bose, S., Broni, S., de L Brooke, M., Butchart, S.H.M., Carlile, N., Catry, P., Catry, T., Charteris, M., Cherel, Y., Clark, B.L., Clay, T.A., Cole, N.C., Conners, M.G., Debski, I., Delord, K., Egevang, C., Elliot, G., Esefeld, J., Facer, C., Fayet, A.L., Fijn, R.C., Fischer, J.H., Franklin, K.A., Gilg, O., Gill, J.A., Granadeiro, J.P., Guilford, T., Handley, J.M., Hanssen, S.A., Hawkes, L.A., Hedd, A., Jaeger, A., Jones, C.G., Jones, C.W., Kopp, M., Krietsch, J., Landers, T.J., Lang, J., Le Corre, M., Mallory, M.L., Masello, J.F., Maxwell, S.M., Medrano, F., Militão, T., Millar, C.D., Moe, B., Montevecchi, W.A., Navarro-Herrero, L., Neves, V.C., Nicholls, D.G., Nicoll, M.A.C., Norris, K., O’Dwyer, T.W., Parker, G.C., Peter, H.U., Phillips, R.A., Quillfeldt, P., Ramos, J.A., Ramos, R., Rayner, M.J., Rexer-Huber, K., Ronconi, R.A., Ruhomaun, K., Ryan, P.G., Sagar, P.M., Saldanha, S., Schmidt, N.M., Schultz, H., Shaffer, S.A., Stenhouse, I.J., Takahashi, A., Tatayah, V., Taylor, G.A., Thompson, D.R., Thompson, T., van Bemmelen, R., Vicente-Sastre, D., Vigfúsdottir, F., Walker, K.J., Watts, J., Weimerskirch, H., Yamamoto, T. & Davies, T.E.; 2025; Global Marine Flyways Identified for Long-Distance Migrating Seabirds From Tracking Data; Global Ecology and Biogeography; 3410.1111/geb.70004.

Oosthuizen, W.C., Bester, M.N., Altwegg, R., McIntyre, T. & de Bruyn, P.J.N.; 2015; Decomposing the variance in southern elephant seal weaning mass: partitioning environmental signals and maternal effects; Ecosphere; 6; art139; 10.1890/ES14-00508.1.

Plaza, P.I., Gamarra-Toledo, V., Rodriguez Eugui, J., Rosciano, N. & Lambertucci, S.A.; 2024; Pacific and Atlantic sea lion mortality caused by highly pathogenic Avian Influenza A(H5N1) in South America; Travel Medicine and Infectious Disease; 59; 102712; 10.1016/j.tmaid.2024.102712.

Puryear, W.B. & Runstadler, J.A.; 2024; High-pathogenicity avian influenza in wildlife: a changing disease dynamic that is expanding in wild birds and having an increasing impact on a growing number of mammals; Journal of the American Veterinary Association; 262; 601–609; 10.2460/javma.24.01.0053.

Riaz, J., Orben, R.A., Gamble, A., Catry, P., Granadeiro, J.P., Campioni, L., Tierney, M. & Baylis, A.M.M.; 2024; Coastal connectivity of marine predators over the Patagonian Shelf during the highly pathogenic avian influenza outbreak; Ecography; 202410.1111/ecog.07415.

Robinson, P.W., Costa, D.P., Crocker, D.E., Gallo-Reynoso, J.P., Champagne, C.D., Fowler, M.A., Goetsch, C., Goetz, K.T., Hassrick, J.L., Hückstädt, L.A., Kuhn, C.E., Maresh, J.L., Maxwell, S.M., McDonald, B.I., Peterson, S.H., Simmons, S.E., Teutschel, N.M., Villegas-Amtmann, S. & Yoda, K.; 2012; Foraging Behavior and Success of a Mesopelagic Predator in the Northeast Pacific Ocean: Insights from a Data-Rich Species, the Northern Elephant Seal; PLoS One; 7; e36728; 10.1371/journal.pone.0036728.

Root, J.J., Porter, S.M., Lenoch, J.B., Ellis, J.W. & Bosco-Lauth, A.M.; 2024; Susceptibilities and viral shedding of peridomestic wildlife infected with clade 2.3.4.4b highly pathogenic avian influenza virus (H5N1); Virology; 600; 110231; 10.1016/j.virol.2024.110231.

Rupprecht, C.E., Bannazadeh Baghi, H., Del Rio Vilas, V.J., Gibson, A.D., Lohr, F., Meslin, F.X., Seetahal, J.F.R., Shervell, K. & Gamble, L.; 2018; Historical, current and expected future occurrence of rabies in enzootic regions; Scientific and Technical Review 37; 729–739; 10.20506/rst.37.2.2836.

Rupprecht, C.E. & Belsare, A.V.; 2026; Rabies and Pinnipeds Reviewed: Premonitions, Perturbations, and Projections?; Veterinary Sciences 1310.3390/vetsci13020200.

Sequeira, A.M.M., Rodriguez, J.P., Marley, S.A., Calich, H.J., van der Mheen, M., VanCompernolle, M., Arrowsmith, L.M., Peel, L.R., Queiroz, N., Vedor, M., da Costa, I., Mucientes, G., Couto, A., Humphries, N.E., Abalo-Morla, S., Abascal, F.J., Abercrombie, D.L., Abrantes, K., Abreu-Grobois, F.A., Afonso, A.S., Afonso, P., Ahonen, H., Akesson, S., Alfaro-Shigueto, J., Andrews, R.D., Angelier, F., Antonopoulou, M., Arata, J.A., Araujo, G., Arauz, R., Arcos, J.M., Arregui, I., Arrizabalaga, H., Auger-Methe, M., Bach, S., Bailleul, F., Baird, R.W., Balazs, G.H., Barco, S.G., Barnett, A., Baverstock, W., Baylis, A.M.M., Beard, A., Becares, J., Belda, E.J., Bell, I., Bennison, A., Benson, S.R., Bernal, D., Berumen, M.L., Bessudo, S., Bezerra, N.P.A., Blaison, A.V., Blanco, G.S., Block, B.A., Bolton, M., Bond, M.E., Bonfil, R., Braun, C.D., Broderick, A.C., Brooke, M.L., Brooks, A.M.L., Brooks, E.J., Bruno, I.M., Burns, J.M., Byrne, M.E., Campana, S.E., Campbell, H.A., Campbell, R.A., Carlisle, A., Carmichael, R.H., Carroll, G., Casale, P., Ceia, F.R., Chapman, D.D., Chapple, T.K., Charrassin, J.B., Chiaradia, A., Chisholm, J., Clarke, C.R., Clay, T.A., Cleguer, C., Clingham, E., Clua, E.E.G., Cochran, J.E.M., Constantine, R., Cooper, R.W., Crochelet, E., Cronin, M., Cuevas, E., DaCosta, K.P., Dagorn, L., Daly, R., Davis, R.W., de Bruyn, P.J.N., Delgado-Trejo, C., Dellinger, T., Derville, S., Diamant, S., DiMatteo, A., Dodge, K.L., Doherty, P.D., Double, M.C., Dove, A.D.M., Doyle, T.K., Drew, M.J., Dubbs, L.L., Duffy, C.A.J., Dutton, P.H., Edwards, E.W.J., Einoder, L.D., Erdmann, M.V., Espinoza, E., Esteban, N., Fagundes, A.I., Feare, C., Ferguson, S.H., Ferreira, L.C., Ferretti, F., Filmalter, J., Finucci, B., Fischer, G.C., Fitzpatrick, R.J., Fontes, J., Formia, A., Fossette, S., Francis, M.P., Friedlaender, A.S., Furtado, M., Gallagher, A.J., Garrigue, C., Gennari, E., Gilchrist, H.G., Godley, B.J., Goldsworthy, S.D., Gollock, M., Carman, V.G., Grecian, W.J., Green, J.R., Guinet, C., Gustafson, J., Guttridge, T.L., Guzman, H.M., Hamer, D., Hamer, K.C., Hammerschlag, N., Hammill, M.O., Harman, L., Harrison, E., Hart, C.E., Harris, A.E., Hastie, G., Hazin, F.H.V., Heard, M., Hearn, A.R., Heide-Jorgensen, M.P., Henry, L., Henry, R.W., 3rd, Hernandez, V.G., Herrera, A.E., Hindell, M.A., Holdsworth, J.C., Holmes, B.J., Howey, L.A., Hoyos Padilla, E.M., Huckstadt, L.A., Hueter, R.E., Lara, P.H., Hussey, N.E., Huveneers, C., Hyland, K., Irion, D.T., Jacoby, D.M.P., Jaeger, A., Jaidah, M.Y., Jessopp, M., Jewell, O.J.D., Johnson, R., Jones, C.G., Jonsen, I.D., Jordan, L.K.B., Jorgensen, S.J., Kato, A., Ketchum, J.T., Kitaysky, A.S., Klimley, A.P., Kock, A.A., Koen, P., Archila, F.L., Lana, F.O., Lane, J.V., Le Corre, M., Lea, M.A., Lea, J., Leat, E.H.K., Lee, O.A., Levenson, J.J., Ley-Quinonez, C.P., Llewellyn, F., Lockhart, G., Lopez, G.G., Mendilaharsu, M.L., Lowther, A.D., Luschi, P., Lutcavage, M.E., Lyon, W.S., Macena, B.C.L., Mackay, A.I., Madden, C.A., Mallory, M.L., Mangel, J.C., Manning, M., Mansfield, K.L., March, D., Marco, A., Marcoux, M., Acuna-Marrero, D., Marsh, H., Marshall, H., Mate, B., McAllister, J.D., McGuire, R.L., McKenzie, J., McLeay, L., McMahon, C.R., Modest, M., Morris, J., Muelbert, M.M.C., Namboothri, N., Nichols, W.J., Nicoll, M.A.C., Norman, B.M., Norris, K., Olsen, E., Oppel, S., Orlowski, S., Pagano, A.M., Page, B., Paiva, V.H., Palacios, D.M., Papastamatiou, Y.P., Parker, D.M., Pattiaratchi, C., Peckham, H., Penaherrera-Palma, C.R., Pepperell, J.G., Phillips, R.A., Pierce, S.J., Pikesley, S.K., Pilcher, N.J., Pinet, P., Pinkerton, M., Pirotta, E., Plot, V., Powell, A.N., Powers, K.D., Prebble, C.E.M., Preston, T.J., Prieto, R., Prosdocimi, L., Quinn, J.L., Quintero, L.M., Raclot, T., Ramirez, I., Ramirez-Macias, D., Ramos, J.A., Read, A.J., Ream, R., Rees, A.F., Reina, R.D., Reisinger, R.R., Revuelta, O., Reynolds, S.D., Richardson, A.J., Riekkola, L., Riet-Sapriza, F.G., Robinson, D.P., Robinson, P.W., Rocha, C.F.D., Rogers, T.L., Rohner, C.A., Ropert-Coudert, Y., Ross, M., Rowat, D.R.L., Ruhomaun, K., Sagar, P.M., Samoilys, M.A., Sanchez, S., Sandoval-Lugo, A.G., Dos Santos, E.A.P., Santos, A.M., Scales, K.L., Schofield, G., Semmens, J.M., Setyawan, E., Shaffer, S.A., Shanker, K., Sheaves, M., Shillinger, G.L., Shivji, M.S., Sianipar, A., Silk, J.R.D., Silva, M.A., Sim, J., Simpson, S.J., Skomal, G., Slip, D.J., Smale, M.J., Soler, G.A., Soria, M., Sousa, L.L., Southall, E.J., Stahl, J.C., Stehfest, K.M., Sterling, J.T., Stevens, J.D., Stevens, G.M.W., Stewart, J.D., Swaminathan, A., Takahashi, A., Tatayah, V., Thiebot, J.B., Thompson, P.M., Thorrold, S.R., Thums, M., Tomas, J., Torres, L.G., Towner, A., Trathan, P.N., Tyminski, J.P., Sagarminaga van Buiten, R., Van Dam, R.P., Vandeperre, F., Varo-Cruz, N., Vaudo, J.J., Vely, M., Villegas-Amtmann, S., Vincent, C., Waayers, D., Wanless, S., Watanabe, Y.Y., Watt, C.A., Weber, S.B., Weber, N., Weise, M.J., Welch, L., Wells, R.S., Werry, J.M., Wetherbee, B.M., White, T.D., Whiting, S.D., Whiting, A.U., Wiebkin, A., Wienecke, B., Wildermann, N.E., Wiley, D.N., Will, A., Williams, S., Windstein, M., Wischnewski, S., Witt, M.J., Womersley, F.C., Wood, A.G., Wright, L.J., Xavier, J.C., Yamamoto, T., Yurkowski, D.J., Zarate, P.M., Zavala-Norzagaray, A., Zerbini, A.N., Costa, D.P., Harcourt, R., Meekan, M.G., Hays, G.C., Sims, D.W., Duarte, C.M. & Eguiluz, V.M.; 2025; Global tracking of marine megafauna space use reveals how to achieve conservation targets; Science; 388; 1086–1097; 10.1126/science.adl0239.

Sobolev, I., Alekseev, A., Sharshov, K., Chistyaeva, M., Ivanov, A., Kurskaya, O., Ohlopkova, O., Moshkin, A., Derko, A., Loginova, A., Solomatina, M., Gadzhiev, A., Bi, Y. & Shestopalov, A.; 2024; Highly Pathogenic Avian Influenza A(H5N1) Virus Clade 2.3.4.4b Infections in Seals, Russia, 2023; Emerging Infectious Diseases; 30; 2160–2164; 10.3201/eid3010.231728.

Stanislawek, W.L., Tana, T., Rawdon, T.G., Cork, S.C., Chen, K., Fatoyinbo, H., Cogger, N., Webby, R.J., Webster, R.G., Joyce, M., Tuboltsev, M.A., Orr, D., Ohneiser, S., Watts, J., Riegen, A.C., McDougall, M., Klee, D. & O’Keefe, J.S.; 2024; Avian influenza viruses in New Zealand wild birds, with an emphasis on subtypes H5 and H7: Their distinctive epidemiology and genomic properties; PLoS One; 19; e0303756; 10.1371/journal.pone.0303756.

Thompson, D.R., Goetz, K.T., Sagar, P.M., Torres, L.G., Kroeger, C.E., Sztukowski, L.A., Orben, R.A., Hoskins, A.J. & Phillips, R.A.; 2021; The year-round distribution and habitat preferences of Campbell albatross (*Thalassarche impavida*); Aquatic Conservation: Marine and Freshwater Ecosystems; 31; 2967–2978; 10.1002/aqc.3685.

Tomas, G., Marandino, A., Panzera, Y., Rodriguez, S., Wallau, G.L., Dezordi, F.Z., Perez, R., Bassetti, L., Negro, R., Williman, J., Uriarte, V., Grazioli, F., Leizagoyen, C., Riveron, S., Coronel, J., Bello, S., Paez, E., Lima, M., Mendez, V. & Perez, R.; 2024; Highly pathogenic avian influenza H5N1 virus infections in pinnipeds and seabirds in Uruguay: Implications for bird-mammal transmission in South America; Virus Evolution; 10; veae031; 10.1093/ve/veae031.

Torres, L.G., Sutton, P.J., Thompson, D.R., Delord, K., Weimerskirch, H., Sagar, P.M., Sommer, E., Dilley, B.J., Ryan, P.G. & Phillips, R.A.; 2015; Poor transferability of species distribution models for a pelagic predator, the grey petrel, indicates contrasting habitat preferences across ocean basins; PLoS One; 10; e0120014; 10.1371/journal.pone.0120014.

Uhart, M. & Vanstreels, R.; 2025; Overview of high pathogenicity avian influenza H5N1 clade 2.3.4.4b in wildlife from Central and South America, October 2022-September 2025; Canadian Journal of Microbiology; 71; 1–8; 10.1139/cjm-2025-0189.

Uhart, M.M., Vanstreels, R.E.T., Nelson, M.I., Olivera, V., Campagna, J., Zavattieri, V., Lemey, P., Campagna, C., Falabella, V. & Rimondi, A.; 2024; Epidemiological data of an influenza A/H5N1 outbreak in elephant seals in Argentina indicates mammal-to-mammal transmission; Nature Communications; 1510.1038/s41467-024-53766-5.

Van Bressem, M.F., Raga, J.A., Di Guardo, G., Jepson, P.D., Duignan, P.J., Siebert, U., Barrett, T., Santos, M.C., Moreno, I.B., Siciliano, S., Aguilar, A. & Van Waerebeek, K.; 2009; Emerging infectious diseases in cetaceans worldwide and the possible role of environmental stressors; Diseases of Aquatic Organisms 86; 143–157; 10.3354/dao02101.

van den Hoff, J., Davies, R. & Burton, H., R,; 2003; Origins, age composition and change in numbers of moulting southern elephant seals (*Mirounga leonina* L.) in the Windmill islands, Vincennes Bay, East Antarctica; Wildlife Research; 30; 275 – 280; 10.1071/WR01086.

Wille, M., Atkinson, R., Barr, I.G., Burgoyne, C., Bond, A.L., Boyle, D., Christie, M., Dewar, M., Douglas, T., Fitzwater, T., Hassell, C., Jessop, R., Klaassen, H., Lavers, J.L., Leung, K.K., Ringma, J., Sutherland, D.R. & Klaassen, M.; 2024a; Long-Distance Avian Migrants Fail to Bring 2.3.4.4b HPAI H5N1 Into Australia for a Second Year in a Row; Influenza and Other Respiratory Viruses; 18; e13281; 10.1111/irv.13281.

Wille, M., Dewar, M.L., Claes, F., Thielen, P. & Karlsson, E.A.; 2024b; A call to innovate Antarctic avian influenza surveillance; Trends in Ecology & Evolution; 10.1016/j.tree.2024.11.005.

Williams, M., Garner, H., Powlesland, R., Robertson, H. & Taylor, G. (2006) Migrations and movements of birds to New Zealand and surrounding seas. pp. 32. Department of Conservation.

Woods, R., Reiss, A., Cox-Witton, K., Grillo, T. & Peters, A.; 2019; The Importance of Wildlife Disease Monitoring as Part of Global Surveillance for Zoonotic Diseases: The Role of Australia; Tropical Medicine and Infectious Disease; 410.3390/tropicalmed4010029.

Xie, R., Edwards, K.M., Wille, M., Wei, X., Wong, S.S., Zanin, M., El-Shesheny, R., Ducatez, M., Poon, L.L.M., Kayali, G., Webby, R.J. & Dhanasekaran, V.; 2023; The episodic resurgence of highly pathogenic avian influenza H5 virus; Nature; 622; 810–817; 10.1038/s41586-023-06631-2.

Xie, Z., Yang, J., Jiao, W., Li, X., Iqbal, M., Liao, M. & Dai, M.; 2025; Clade 2.3.4.4b highly pathogenic avian influenza H5N1 viruses: knowns, unknowns, and challenges; Journal of Virology; 99; e0042425; 10.1128/jvi.00424-25.

Younger, J.L., Patier, L., Brav-Cubitt, T., van Tonder, A., Clessin, A., Coleman, J., Farrer, J., McMahon, C.R., Gamble, A. & Fountain-Jones, N.; 2026; High pathogenicity avian influenza virus H5N1 clade 2.3.4.4b in Antarctica: Multiple Introductions and the First Confirmed Infection of Ice-Dependent Seals; bioRxiv; 10.64898/2026.01.04.697571.

